# Regional gene expression patterns are associated with task-specific brain activation during reward and emotion processing measured with functional MRI

**DOI:** 10.1101/2020.06.27.175257

**Authors:** Arkadiusz Komorowski, Ramon Vidal, Aditya Singh, Matej Murgaš, Tonatiuh Pena-Centeno, Gregor Gryglewski, Siegfried Kasper, Jens Wiltfang, Rupert Lanzenberger, Roberto Goya-Maldonado

## Abstract

The spatial relationship between gene expression profiles and neural response patterns known to be altered in neuropsychiatric disorders, e.g. depression, can guide the development of more targeted therapies. Here, we estimated the correlation between human transcriptome data and two different brain activation maps measured with functional magnetic resonance imaging (fMRI) in healthy subjects. Whole-brain activation patterns evoked during an emotional face recognition task were associated with topological mRNA expression of genes involved in cellular transport. In contrast, fMRI activation patterns related to the acceptance of monetary rewards were associated with genes implicated in neuronal development, synapse regulation, and gene transcription. An overlap of these genes with risk genes from major depressive disorder genome-wide associations studies revealed the involvement of the master regulators TCF4 and MEF2C. These results were further validated by means of meta-analytic fMRI data. Overall, the identification of stable relationships between spatial gene expression profiles and fMRI data may reshape the prospects for imaging transcriptomics studies.

## 1 Introduction

Over the last decades, genetic and neuroimaging studies have significantly contributed to current knowledge about human neural functions. While individual neuroscientific methods facilitated the comprehension of physiological processes as well as pathological alterations in psychiatric disorders, a multimodal integration of large-scale data has proven to be even more conducive for in-depth understanding (Kitchen et al., 2014; Medland et al., 2014). For example, neural correlates of common psychological processes can be quantified by functional magnetic resonance imaging (fMRI) with high spatial resolution. The fact that the measured signal patterns are to some extend susceptible to genotype variations (Rose & Donohoe, 2013; Wolf et al., 2015), however, illustrates the conceptual challenges of studies linking data across disparate scales of biological resolution. Albeit investigating the influence of the genome on pre-selected quantitative traits, imaging genetics studies mostly ignore transcriptional regulation. On the contrary, post-mortem gene expression data from the Allen Human Brain Atlas (AHBA) can be applied to investigate the relationship between the transcriptome and protein distribution (Komorowski et al., 2017), brain morphology (Shin et al., 2018), or connectivity (Richiardi et al., 2015). Recent studies even integrated characteristic imaging findings to assess regional gene expression profiles of individual genes within the scope of neurological and psychiatric disorders (Freeze et al., 2018; Romme et al., 2017). Yet, in the light of the complexity of the human brain it appears beneficial to include the whole transcriptome for the evaluation of potential influences on neuroimaging properties. It can be argued that numerous differentially expressed genes of varying significance affect functional brain activation during specific psychological processes. Although a promising approach to contrast fMRI activation and the transcriptome making use of the meta-analytic Neurosynth database was presented by Fox et al. (2014), their study suffered from a low spatial resolution and also lacked disease-related aspects. Furthermore, the impact of master regulators implicated in neuropsychiatric disorders on the expression of genes that might influence in vivo regional blood oxygenation level dependent (BOLD) signaling has not been resolved yet.

From all psychiatric disorders, major depressive disorder (MDD) is now the leading cause of disability worldwide (James et al., 2018). MDD strongly contributes to the overall global burden of disease with an increasing prevalence over the years, whereby additive genetic effects attribute to approximately 9 % of the variation in liability of this disorder (Wray et al., 2018). Notably, the differential expression of genes between depressed individuals and the general population highlights the relevance of specific transcriptomic signatures for human brain function (Ciobanu et al., 2016; Mehta et al., 2010). In this regard, paradigms examining prominent behavioral elements such as impaired affect modulation or loss of interest and pleasure in common experiences are amongst the best-established in neuroimaging studies, which justifies their application to investigate core depressive symptoms (Foland-Ross & Gotlib, 2012). Alterations of BOLD reactivity during processing of negatively valenced information or incentive-based learning thereby drive the conceptualization of major domains of functioning within the realms of the Research Domain Criteria (RDoC) framework (Sanislow et al., 2019), spanning from a physiological to a more critical pathological state. Since brain activation patterns during emotion and reward processing are particularly responsive to depressive symptoms, a major role of MDD risk genes is assumed for both imaging paradigms.

The goal of this study was to evaluate the spatial relationship between whole-transcriptome expression maps and specific brain activation patterns measured in healthy human subjects during emotion and reward processing in order to reveal associations of task-based fMRI data with biological processes according to the Gene Ontology (GO) database. Ultimately, we aimed to identify master regulating genes implicated in depressive disorders that exert potential effects on the investigated imaging paradigms.

## 2 Materials and Methods

### 2.1 Participants

Healthy subjects were recruited from the university environment and gave written informed consent to the study procedures previously approved by the Ethics Committee of the University Medical Center Göttingen. Included participants (aged between 22 and 52 years; M = 40.5, SD = 14.37) were of Caucasian European ethnicity and fluent in German language. Exclusion criteria comprised contraindications to MRI, past or present psychiatric, neurological, or medical disorders, consumption of psychotropic drugs, and a positive family history of psychiatric disorders. In total, 26 men and 22 women completed two fMRI paradigms related to emotional face recognition and reward processing. Excessive movement in any of the three translation (> 2 mm) or rotation (> 2°) planes resulted in exclusion of 4 participants.

### 2.2 Functional brain imaging

Functional imaging data was acquired using a 3 T scanner (Siemens Magnetom TRIO, Siemens Healthcare, Erlangen, Germany) and a 32-channel head coil with a 2 x 2 x 2 mm voxel size, TR 2500 ms, TE 33 ms, 70° flip angle, 10 % distance factor, FOV 256 mm and 60 slices with multiband factor of 3 for the acquisition of T2*-weighted images. Imaging data analysis was performed using Statistical Parameter Mapping (SPM12; Wellcome Department of Imaging Neuroscience, Institute of Neurology, London, UK) and Matlab R2015b (The Mathworks Inc., Natick, MA, USA). First, echo planar imaging (EPI) images were standardly preprocessed with slice time correction, realignment, and normalization into the Montreal Neurological Institute (MNI) space, as well as smoothing with an 8 x 8 x 8 mm FWHM Gaussian kernel.

Null hypotheses relating to random fMRI activation during different psychological states were tested for both imaging paradigms. First, specific activation maps reflected brain activation during performance of the tasks. In contrast, corresponding control conditions represented non-specific hemodynamic activity inherent to any task performance during fMRI measurement, caused by unspecific physiological activation, e.g. related to visual, auditory, attentional or motor functions. Estimates of neural activity were initially computed with a general linear model (GLM) for each subject individually (first-level analysis) with nuisance movement parameters regressed as covariates-of-no-interest. Later, experimental and control conditions were evaluated at group level (second-level analysis) and resulting activation maps, which represented task-specific brain activation, were used for further analyses.

### 2.3 Emotional face recognition

The paradigm of implicit emotional face recognition contains two different contexts: human faces and geometric objects. Pictures of males and females with varying negative face expressions obtained from the Radboud database (Langner et al., 2010) were presented for 17 s, during which participants responded to the gender of the presented person with a button press. Thereby, perception of emotions was rather implicit, which has been shown to enhance the activation of emotional correlates (Keightley et al., 2003). For the control condition, participants were instructed to respond analogously to the shape of an object, either an ellipse or a rectangle, positioned in the face area and made from scrambling original face trials. All trials were controlled for brightness, contrast and presented in a very similar composition. The activation patterns representing the experimental and control conditions were computed using first-level (single-subject) contrasts of the trials from emotional faces and object blocks, respectively. Resulting data were then used for second-level (group) analysis, as standardly performed for random effects model.

### 2.4 Reward processing

For this study, a previously established fMRI paradigm was implemented, which has been broadly used to investigate physiological and pathological reward mechanisms (Diekhof & Gruber, 2010; Goya-Maldonado et al., 2015). Briefly, participants performed a modified delayed match to sample task, including two contexts involving previously conditioned stimuli to monetary rewards: acceptance or rejection of rewards, i.e. pressing a button when squares are shown. Subjects were instructed they would receive 30 € for their participation, and that they were able to double this amount according to their task performance. As control trials, subjects responded with a button press to stimuli that required motor performance as well as attentional and memory resources, but were not conditioned to monetary reward. To compute the control condition, first-level (single-subject) contrasts of correctly matched sample trials within the same experimental block of reward trials were used. For experimental conditions, first-level experimental contrasts were calculated from brain activation elicited during acceptance of previously conditioned stimuli. At group-level, activations related to experimental and control trials were contrasted to obtain functional activation patterns.

### 2.5 Meta-analytic functional brain activation

Besides fMRI data obtained from participants performing two different tasks at our institution, we evaluated large-scale meta-analytical imaging data from the Neurosynth platform (https://neurosynth.org/), which provides probabilistic brain activation maps computed from an automated meta-analysis based on published fMRI studies. This online database combines text-mining and machine-learning techniques to generate statistical inference maps of currently 1,335 imaging terms from 14,371 fMRI studies including male and female participants (Yarkoni et al., 2011). Within the framework of the Neurosynth database, particular psychological processes are labeled with terms of interest and represented by uniformity test maps. For this study, activation maps related to emotion (“fearful faces”) and reward (“rewards”) processing were downloaded in MNI152 space with 2 mm resolution to validate fMRI data obtained at our institution. Each uniformity test map provides z-scores, which reflect the proportion of studies that report cerebral activation at a given voxel. Thus, obtained Neurosynth maps depicted specific brain areas that were consistently reported in studies investigating fMRI activation for emotion and reward processing, respectively. In a first step, activation maps, with z-scores representing how brain regions are related to the chosen term, were generated from the Neurosynth online database. Since the online user interface only provides thresholded maps, we used the Neurosynth toolbox (https://github.com/neurosynth/neurosynth) for python to download unthresholded maps that were further smoothed using 8 x 8 x 8 mm FWHM to match the kernel size of single-site data. Assessment of conformity between meta-analytic data and measured fMRI maps was performed qualitatively and quantitatively, whereby region-wise Spearman’s correlation coefficients were calculated in cortical and subcortical regions. The Neurosynth terms “fearful faces” and “rewards” matched well with functional activation related to recognition of negative faces and the acceptance of monetary rewards, respectively (Supplementary Fig. 1A, B). Based on low fMRI activity, cerebellum was not considered for the comparison. In contrast to single-site fMRI data, meta-analytical information comprises rather positive values due to the sparse reporting of brain regions showing negative activation in most neuroimaging studies. Hence, when processing unthresholded data from the uniformity test maps, mainly positive values determined the association analysis between single-site and Neurosynth data.

### 2.6 Whole-brain gene expression

The AHBA (www.brain-map.org) consists of microarray assessments from 3,702 brain tissue samples collected across 6 human donors (1 female, mean age = 42.5, SD = 13.4) derived from diverse regions of the brain, extensively described in the original publications (Hawrylycz et al., 2012; Shen et al., 2012). As delineated by other authors (Arnatkevičiūtė et al., 2019), multiple data processing steps such as gene annotation, probe selection, or sample assignment need to be considered to facilitate correlation analyses between gene expression and neuroimaging data. By using common parcellation schemes, expression levels in numerous brain regions devoid of tissue samples remain indefinite, which necessitates an interpolation of the AHBA data to meet the resolution of neuroimaging techniques. Therefore, high-resolution human transcriptome maps were obtained from a publicly accessible database (available for download at www.meduniwien.ac.at/neuroimaging/mRNA.html) for all analyses of this study. The methodology regarding the creation of these maps was described extensively in the original publication (Gryglewski et al., 2018). Briefly, seamless gene expression values (log2) were predicted by the authors across the whole brain to compensate for the sparse anatomical sampling of the AHBA. Using Gaussian process regression, missing expression values were inferred by Gryglewski et al. (2018) at all cortical, subcortical, and cerebellar structures for 18,686 genes. While normalization for inter-individual differences between mRNA probes and donor brains attributable to high efforts in tissue preparation was performed, sex-specific differences were not considered due to the low validity of female transcriptome data. The interpolated expression maps were associated with Entrez Gene IDs and registered to MNI space in line with common neuroimaging techniques, except for insensitive probes or missing allocations to gene IDs.

### 2.7 Spatial correlation between gene expression and brain activity

Initially, gene lists ranking correlations between transcriptome maps and functional brain activation patterns elicited by selected paradigms were compiled, whereby the ranking of each gene depended on its correlation strength with the corresponding single-site BOLD activation map. After initial inspection, the cerebellum was excluded from further analysis due to marginal activation during both fMRI paradigms. On the basis of known differences in gene expression between broad anatomical regions (Chen et al., 2016; Hawrylycz et al., 2012), correlation analyses were assessed within cortical and subcortical regions separately (Supplementary Fig. 2A, B). All available transcriptome maps were aligned with group-averaged fMRI activation maps for each paradigm in order to conduct association analyses. To account for partly non-symmetrical distribution of mRNA data and existence of outliers, Spearman’s correlation coefficients were calculated between each gene-imaging pair (mRNA expression vs. fMRI activation). Main findings are reported for region-wise analyses, along with additional results for voxel-wise correlations (total number of voxels was 129,817 in the cortex and 10,863 in subcortex; zero-values outside of the investigated area were excluded). Each statistical map was segmented into brain regions according to the Brainnetome atlas, which was selected for primary analyses, because it labels a sufficient number of subcortical (n = 36) and cortical (n = 210) regions-of-interest (ROIs) (Fan et al., 2016). A complementary analysis with fewer brain regions (12 subcortical and 78 cortical ROIs) was done using the automated anatomical labeling (AAL) brain atlas (Tzourio-Mazoyer et al., 2002) to evaluate influences of different parcellation methods. Both atlases were aligned with fMRI data and the transcriptome maps in MNI space using SPM12, while extraction of ROIs and correlation analyses were performed in MATLAB2018a (www.mathworks.com). For each task, the null model was accounted for in the fMRI data analysis yielding specific activation maps related to emotion and reward processing. Separate correlation analyses for the task and control conditions were considered redundant and true associations between transcriptome and imaging maps were assumed for all genes that yielded higher correlation coefficients than would be expected under a random distribution of gene expression data. Thereby, statistical significances of region-wise correlations were assessed by means of randomization tests including 10,000 iterations. For each sampled permutation, mRNA values were randomly shuffled and correlated with fMRI data. Two-sided p-values were calculated as the proportion of sampled permutations, where the absolute value was greater than the true correlation coefficient (non-shuffled data).

### 2.8 Identification of overlap between analyzed data sets

To compare various sets of mRNA-fMRI correlations we used Rank-Rank Hypergeometric Overlap (RRHO) package (version 1.26.0) in R (https://www.bioconductor.org/packages/release/bioc/html/RRHO.html), which allows statistical testing of the extent of overlap between two ranked lists. RRHO determines the degree of differential expression observed in profiling experiments using the hypergeometric distribution. While originally applied for the comparison of gene expression profiles between different microarray platforms or types of model system, we used RRHO to compare genes that ranked according to relevant measures of differential information (in this case the correlation strength with fMRI data). Thereby, genes of two datasets were ranked according to their names and the corresponding ranks were tested for statistical overlap. We provide both a graphical representation of the characteristics of analyzed datasets (corresponding p-values) as well as a statistical measure of overlap (rho_RRHO_). A high overlap implies that positively correlating genes of a given list show high ranks in the second list, while genes with a negative correlation are also negatively associated in the alternative list. Applying this method offered the advantage of using the whole continuum of previously ranked genes without the need to truncate the list by pre-defined thresholds.

### 2.9 Analysis of biological processes

Making use of the GO knowledgebase (Ashburner et al., 2000; The Gene Ontology Consortium, 2018), a comprehensive resource for computational analysis of large-scale data, we explored enriched biological process terms that included genes with expression patterns highly correlated with each fMRI paradigm. Given the disparate biological resolutions of fMRI and gene expression data, the other two GO categories referring to molecular functions and cellular components were not included in this study. Consisting of multiple ordered molecular functions, a biological process term can be differentiated into various levels of specificity. Hereof, only specialized biological processes (GO levels higher than 4) were analyzed to attain conclusive information about underlying genetic substrates and their functions within the emotion and reward systems. GO analysis was performed separately for positively and negatively correlated genes with cut-off values for correlation coefficients above rho = 0.5 and below rho = −0.5, which excluded non-spatially depending genes showing insufficient associations with fMRI data. The Cytoscape plugin “ClueGO” (Bindea et al., 2009) was used to compute GO enrichment (default parameters), comprising all precompiled annotations that were listed at the time of analysis (18,361 biological process terms). For each paradigm, ClueGO generated a binary gene-term matrix based on previously ranked correlation coefficients of the associated genes. P-values were obtained for enriched biological processes by means of hypergeometric tests and subsequently adjusted for multiple testing by applying Bonferroni correction (p_cor_). Exclusively region-wise correlations were considered for the analyses due to more refined results. Absolute numbers and percentages of genes that were statistically overrepresented within each resulting biological process term were calculated. Considering a potential gene overlap between the enriched GO terms, the Jaccard index was used to assess similarity across imaging paradigms.

### 2.10 Association of risk genes implicated in major depression with functional imaging data

Regarding genetic risks for depressive disorders, recently 42 functional and 27 non-functional risk genes implicated in major depression were identified in a genome-wide association meta-analysis by Wray et al. (2018), which included 135,458 cases and 344,901 controls. For each fMRI paradigm, the presence of master regulators among the 42 functional risk genes proposed by Wray et al. (2018) was evaluated (Supplementary Table 1). The analysis was performed by means of the Cytoscape plugin iRegulon, which reliably detects upstream regulators and corresponding master regulons on diverse types of gene sets (Janky et al., 2014). In this study, master regulating genes and their direct transcriptional targets were identified using motif discovery within each set of correlated genes based on previously ranked lists including HGNC symbols. Resulting master regulators for each predicted regulon of the input gene sets were eventually compared with the genes implicated in major depression. All computations were performed separately for positive and negative mRNA-fMRI correlations above rho = 0.6 and below rho = −0.6, respectively. These thresholds were set to ensure a sufficient number of evaluated genes in subcortical regions. In the cortex, insufficient data availability hampered analyses of co-expressed genes due to generally lower correlation coefficients of mRNA-fMRI associations compared to the subcortex. Parameters such as enrichment score and significances were used as default; distance from TSS was set to 500bp.

Additionally, potential enrichment of the risk genes implicated in major depression was assessed for each fMRI paradigm by means of gene set enrichment analysis (GSEA) (Mootha et al., 2003; Subramanian et al., 2005). In terms of the methodology, the GSEA enrichment score (ES) reflects the degree to which an analyzed gene set is overrepresented within a ranked list. Corresponding to a weighted Kolmogorov-Smirnov-like statistic, the ES is calculated by a stepwise increase or decrease of the total sum statistic of the list as described by Subramanian et al. (2005). Here, the implementation in R that is available in the package clusterProfiler was utilized for the enrichment analysis (Yu et al., 2012). Based on initially compiled gene lists that were ranked by correlation strengths with fMRI data, the position of each MDD risk gene was compared to the position of all other genes. We tested, whether the risk genes were randomly distributed throughout each ranked list or primarily found at the top (showing positive correlations with fMRI data) and bottom (showing negative correlations). Ranked correlation coefficients and their corresponding p-values were defined as input factors for the GSEA, which required a summarized biological value for each included gene. Risk genes ranked in higher positions contributed more to the resulting significance of the enrichment score than lower ranked genes. The maximum ES (with positive or negative values) represented the maximum deviation from zero, whereby statistical significance was tested against an ES referring to a null distribution of permuted data. Since enrichment was evaluated for only one gene set, adjustment of the estimated significance level (p < 0.05) for multiple hypothesis testing was dispensable.

## 3 Results

### 3.1 Topological specificity of emotion and reward processing

In conjunction with known abnormalities in social interaction and reward responsiveness of patients with depressive disorders, fMRI activation within the social processes and positive valence systems domains of the RDoC framework was evaluated. We minimized unspecific signal variations related to visual, auditory, attentional, and executive processing by contrasting brain activation elicited by the experimental condition with task control conditions in each participant. Sequentially, second level analyses provided specific activation patterns elicited by emotion and reward processing (full acquisition and analysis pipeline described in Methods section). The applied emotional face recognition task didn’t focus attention on the emotional content of presented faces, but requested gender discrimination instead, provoking rather implicit emotional processing in limbic as well as non-limbic areas such as prefrontal cortices. When testing reward responsiveness, significant activations in dopaminergic brain regions were observed after acceptance of priorly conditioned stimuli, particularly in the mesolimbic reward system (Supplementary Table 2).

To expand the scope of results generated with single-site fMRI data, we obtained corresponding uniformity maps from the Neurosynth database that inform about the consistency of functional brain activation for particular processes of emotion and reward circuits. Spatial activation clusters detected at our institution were validated using matching statistical inference maps derived from 91 and 246 studies associated with the terms “fearful faces” and “rewards”, respectively. Both meta-analytic maps, which can be obtained online from the Neurosynth framework, were representative of the expected neural correlates elicited by emotional face and reward tasks.

### 3.2 Associations between the transcriptome and functional brain activation

For each paradigm, correlation analyses yielded spatial associations between functional brain activation and mRNA expression for 18,686 individual genes (rankings of genes as well as corresponding Spearman’s correlation coefficients for both datasets are reported in Supplementary Table 3). All correlation analyses were performed separately for cortical and subcortical regions to prevent a possible bias that might arise from gene expression differences between broad anatomical areas. Resulting correlations were highly specific for each paradigm, accounted for by the weak overlap of compiled correlation lists between the two psychological processes, i.e. emotion and reward processing (subcortex: rho_RRHO_ = −0.282; cortex: rho_RRHO_ = 0.205) (Supplementary Fig. 3A, B). In contrast, high agreement of ranked mRNA-fMRI correlations between both applied brain parcellation atlases was observed for emotional face recognition (subcortex: rho_RRHO_ = 0.697; cortex: rho_RRHO_ = 0.822) and acceptance of monetary rewards (subcortex: rho_RRHO_ = 0.748; cortex: rho_RRHO_ = 0.918) (Fig. 1A-C).

**Fig. 1:**
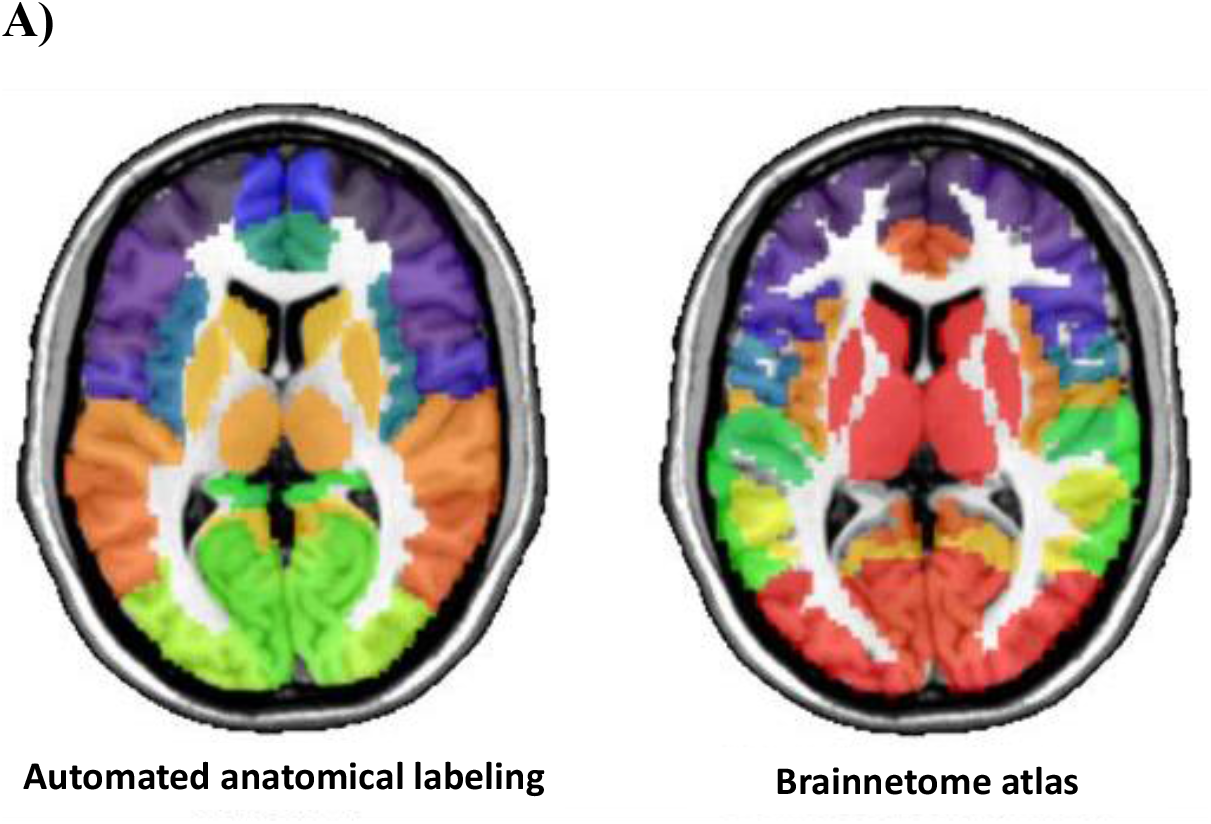
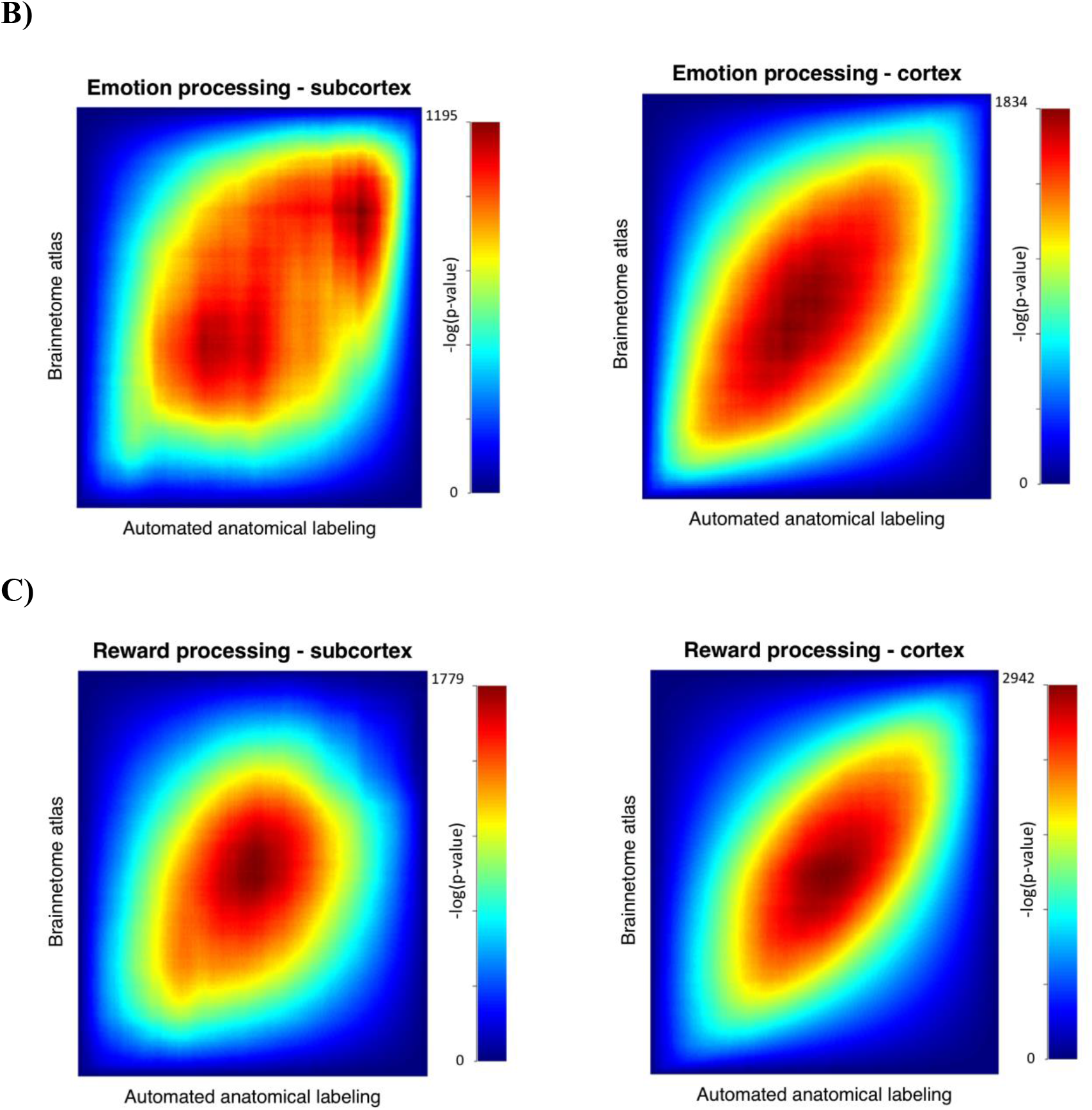
Comparison of single-site imaging data applying two different parcellation schemes. **A)** Visualization of cortical and subcortical regions of interest according to automated anatomical labeling (left) and the Brainnetome atlas (right) (transversal plane in MNI standard space; z = 8). **B)** Region-wise Rank–rank hypergeometric overlap (RRHO) comparing ranked lists including 18,686 genes indicated high agreement between both atlases for emotional face recognition in the subcortex (rho_RRHO_ = 0.697) and cortex (rho_RRHO_ = 0.822). Genes with congruent correlation coefficients (either positive or negative) show higher statistical significance in the bottom left and top right corner. **C)** Likewise, RRHO of both parcellation methods was performed for reward processing in subcortical (rho_RRHO_ = 0.748) and cortical regions (rho_RRHO_ = 0.918).

Regarding emotion processing, single-site brain activity patterns correlated positively as well as negatively with whole-brain transcriptome maps (Fig. 2A, B). Significant region-wise correlations yielded similar results compared to voxel-wise analyses, ranging from rho = −0.739 to rho = 0.865 for subcortical and from rho = −0.449 to rho = 0.431 for cortical regions (Supplementary Fig. 4A, B). Out of 18,686 spatial associations between gene expression and brain function the 15 highest positive correlating genes are listed in Table 1 (p < 0.0001). In subcortical regions, MALL showed the strongest voxel-wise correlation (rho = 0.633), while C10orf125 showed the highest region-wise correlation (rho = 0.865). In the cortex, SPDYA yielded strongest voxel-wise (rho = 0.328) and FOXN4 strongest region-wise (rho = 0.431) correlation (Fig. 2C).

**Fig. 2:**
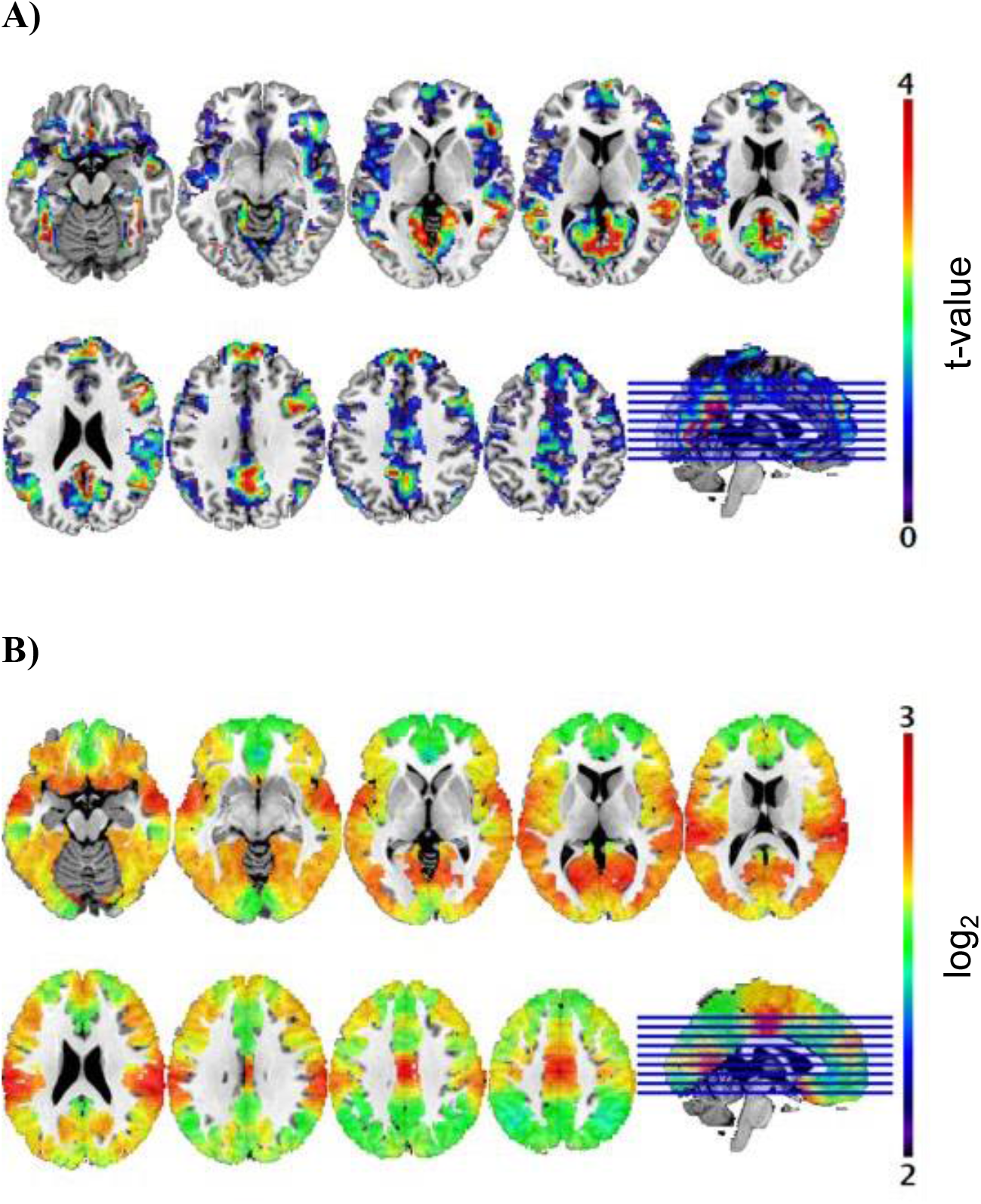
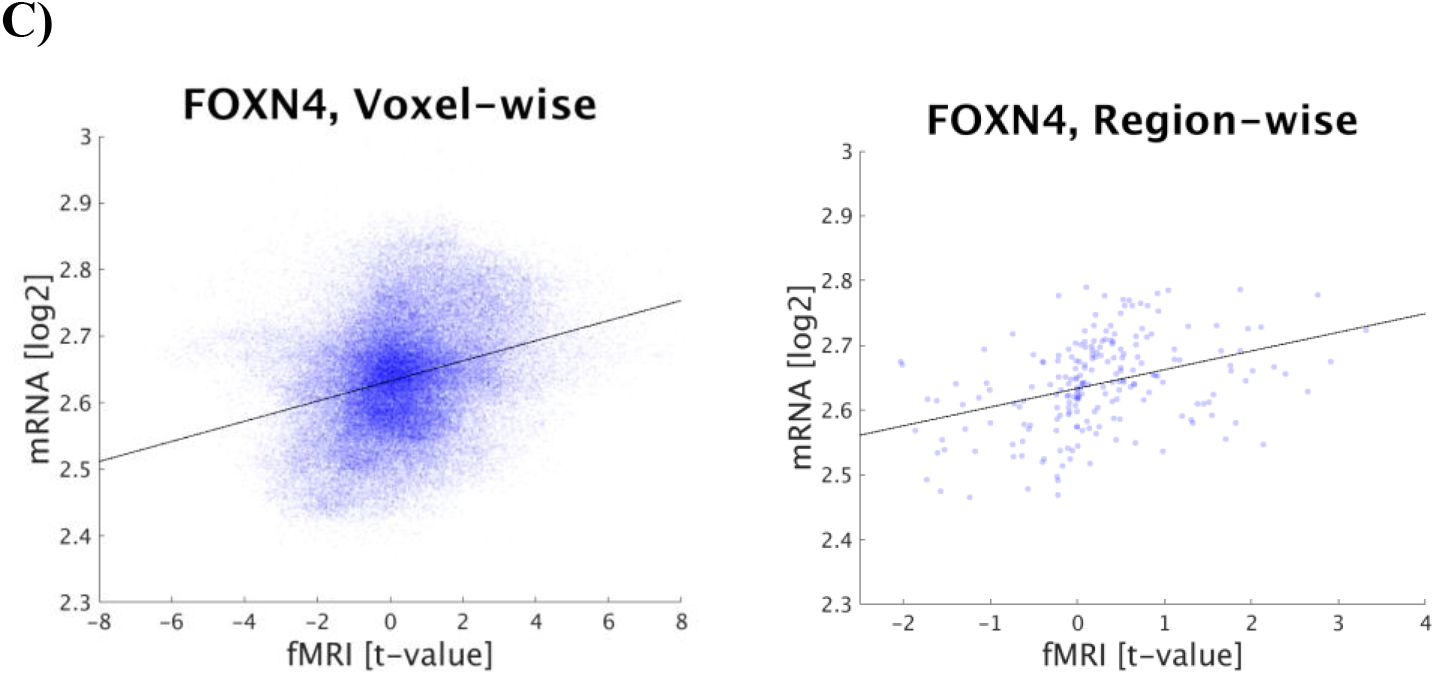
Comparison of task-specific functional brain activation and mRNA expression in cortical regions. **A)** Single-site functional magnetic resonance imaging data (t-value) during emotion processing is visualized in MNI space (activation maps are thresholded above 0 for visualization purposes only). **B)** FOXN4 gene expression is based on cortical transcriptome maps (log_2_) by Gryglewski et al. (2018). **C)** The scatter plots depict correlations between cortical mRNA levels of FOXN4 and imaging data (emotional face recognition) for voxel-wise (rho = 0.285; 129,817 voxels) and region-wise (rho = 0.431; 210 regions, p < 0.0001) analyses. Each dot represents expression values and corresponding imaging parameters at target coordinates or within anatomical regions, respectively.

**Table 1:** Ranking of Spearman’s correlation coefficients for genes with expression patterns showing highest positive associations with single-site imaging data (emotional face processing).

| Subcortex |  |  |  | Cortex |  |  |  |
| --- | --- | --- | --- | --- | --- | --- | --- |
| Voxel-wise correlations |  | Region-wise correlations |  | Voxel-wise correlations |  | Region-wise correlations |  |
| rho | Gene name | rho | Gene name | rho | Gene name | rho | Gene name |
| **0.633** | **MALL** | 0.865 | C10orf125 | 0.328 | SPDYA | **0.431** | **FOXN4** |
| 0.616 | HRASLS5 | 0.845 | PTRH1 | **0.298** | **CCDC62** | 0.395 | PIK3R6 |
| 0.614 | FAT4 | 0.818 | GHRLOS | 0.296 | PYGO2 | 0.390 | RYBP |
| 0.612 | SCARA5 | 0.817 | SLC24A4 | 0.289 | CPZ | 0.387 | STC1 |
| 0.610 | MESP1 | **0.817** | **AC022098.3** | **0.285** | **FOXN4** | **0.384** | **CCDC62** |
| 0.608 | LINC00260 | 0.812 | ZNF280C | 0.283 | FRMD3 | 0.379 | FUBP1 |
| 0.604 | SKAP2 | 0.810 | FUT1 | 0.282 | PHOX2B | 0.378 | SMYD1 |
| 0.602 | SCPEP1 | 0.804 | NLE1 | **0.280** | **ATXN10** | 0.356 | MIA2 |
| 0.596 | CRHBP | 0.803 | FBP1 | 0.276 | XAGE3 | 0.356 | TMPRSS4 |
| 0.594 | RAB3GAP1 | 0.803 | C16orf55 | 0.273 | EGFL6 | 0.350 | DLG3 |
| 0.592 | POLB | **0.800** | **MALL** | 0.273 | KIAA1328 | 0.346 | HSPA1A |
| 0.591 | LOC642852 | 0.798 | PDK1 | 0.272 | RNF215 | 0.345 | CSTA |
| 0.590 | KIAA0947 | 0.798 | B4GALT5 | 0.272 | ATP6V1D | 0.342 | NR4A2 |
| 0.589 | SNAP29 | 0.793 | CCDC105 | 0.272 | ERC2-IT1 | **0.341** | **ATXN10** |
| **0.585** | **AC022098.3** | 0.793 | CAMLG | 0.272 | LINC00158 | 0.335 | AC008026.2 |

Footnote: All listed region-wise correlation coefficients were significant in permutation tests ( $p < 0.0001$ ). Genes marked in bold ranked within the 15 highest positively correlating genes in both voxel-wise and region-wise analyses.

Analogous to the emotion task, ranked lists with gene expression patterns spatially associated with measured imaging data were compiled for the reward system, whereby the 15 highest positive correlating genes are listed in Table 2 (p < 0.0001). Region-wise analyses yielded higher correlation coefficients than the voxel-wise approach with less prominent associations in the cortex (rho = −0.639 to rho = 0.698), compared to subcortical regions (rho = −0.788 to rho = 0.81). In the subcortex, MDK showed the strongest voxel-wise correlation (rho = 0.488) of all 18,686 genes and also a high region-wise correlation coefficient (rho = 0.803) (Fig. 3A–C). Comparing strongest voxel-wise vs. region-wise correlations in the cortex, 11 out of 15 genes were congruent (DUSP3, CA10, PIK3CD, HDAC9, LASS6, GRB14, OLFM3, SHC1, NT5DC2, ASS1, and SPRN), indicating high agreement between both approaches (Supplementary Fig. 5A, B). Notably, gene expression of DUSP3 yielded strongest cortical correlations with reward processing both in the voxel-wise (rho = 0.548) as well as the region-wise analysis (rho = 0.698) (Supplementary Fig. 6).

**Table 2:** Ranking of Spearman’s correlation coefficients for genes with expression patterns showing highest positive associations with single-site imaging data (acceptance of monetary rewards).

| Subcortex |  |  |  | Cortex |  |  |  |
| --- | --- | --- | --- | --- | --- | --- | --- |
| Voxel-wise correlations |  | Region-wise correlations |  | Voxel-wise correlations |  | Region-wise correlations |  |
| $\rho$ | Gene name | $\rho$ | Gene name | $\rho$ | Gene name | $\rho$ | Gene name |
| **0.488** | **MDK** | 0.810 | VMO1 | **0.548** | **DUSP3** | **0.698** | **DUSP3** |
| 0.481 | HELLS | 0.805 | OSTM1 | **0.547** | **CA10** | **0.681** | **CA10** |
| 0.475 | RBBP8 | **0.803** | **MDK** | **0.542** | **PIK3CD** | **0.680** | **PIK3CD** |
| **0.453** | **ATF1** | **0.798** | **KRT18P19** | **0.533** | **GRB14** | **0.653** | **HDAC9** |
| **0.451** | **KRT18P19** | 0.791 | IMPACT | **0.517** | **ASS1** | **0.640** | **LASS6** |
| 0.450 | CD274 | **0.787** | **USP24** | **0.516** | **LASS6** | 0.639 | CCNYL1 |
| 0.446 | C8orf22 | 0.784 | NEK1 | **0.505** | **HDAC9** | **0.637** | **GRB14** |
| 0.442 | SALL4 | 0.782 | RCBTB2 | 0.504 | FBXL2 | **0.627** | **OLFM3** |
| **0.440** | **USP24** | 0.782 | PCBD2 | **0.497** | **OLFM3** | **0.609** | **SHC1** |
| 0.439 | SFRP5 | **0.781** | **CD99** | 0.492 | TMEM150C | **0.607** | **NT5DC2** |
| 0.438 | PMCH | 0.774 | TRIM34 | **0.477** | **SHC1** | 0.607 | GOLPH3L |
| **0.436** | **CD99** | 0.764 | ARF4 | **0.475** | **SPRN** | **0.606** | **ASS1** |
| 0.435 | C12ORF75 | **0.760** | **ATF1** | **0.473** | **NT5DC2** | 0.601 | MYBPC2 |
| 0.431 | NPB | 0.760 | ADAL | 0.469 | MUM1L1 | 0.595 | MPO |
| 0.430 | RABL3 | 0.758 | HERC5 | 0.463 | ARHGAP8 | **0.592** | **SPRN** |

Footnote: All listed region-wise correlation coefficients were significant in permutation tests ( $p < 0.0001$ ). Genes marked in bold ranked within the 15 highest positively correlating genes in both voxel-wise and region-wise analyses.

**Fig. 3:**
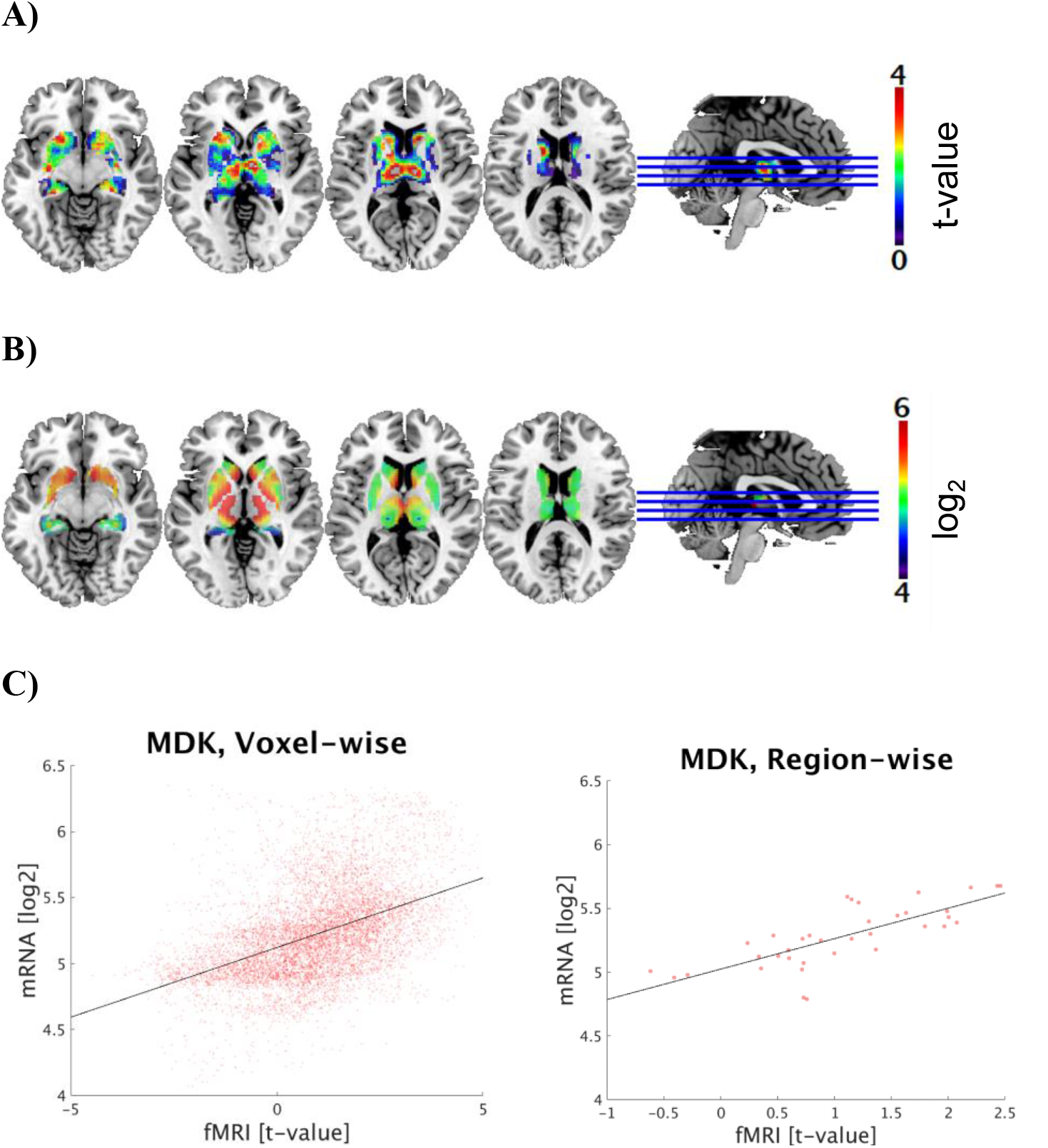
Comparison of task-specific functional brain activation and mRNA expression in subcortical regions. **A)** Single-site functional magnetic resonance imaging data (t-value) during reward processing is visualized in MNI space (activation maps are thresholded above 0 for visualization purposes only). **B)** MDK gene expression is based on subcortical transcriptome maps (log_2_) by Gryglewski et al. (2018). **C)** The scatter plots depict correlations between subcortical mRNA levels of MDK and imaging data (acceptance of monetary rewards) for voxel-wise (rho = 0.488; 10,863 voxels) and region-wise (rho = 0.803; 36 regions, p < 0.0001) analyses. Each dot represents expression values and corresponding imaging parameters at target coordinates or within anatomical regions, respectively.

### 3.3 Ontological analysis of task-specific biological processes

Subsequent analyses of previously compiled mRNA-fMRI correlations revealed multiple genes that were included within gene sets listed in the GO knowledgebase. Thereby, individual GO terms were associated with genes that correlated either positively or negatively with fMRI data after applying Bonferroni correction (p_corr_ < 0.05), indicating a unique relationship between specific biological processes and each task (Fig. 4; Supplementary Table 4). Considering a potential overlap of significantly associated GO terms between both imaging paradigms, rather low Jaccard coefficients indicated distinctive associations of biological processes with different psychological processes (Supplementary Fig. 7). Regarding neural responses during emotional face recognition in subcortical regions, significantly associated biological programs were related to cellular transport processes and mainly included genes that showed positive correlations between gene expression and imaging data. In line with the assumed relevance of molecular transduction for emotion regulation processes, the most significant overlap was observed for the GO term peptide transport. In total, 203 genes positively correlated with emotion processing were also present within this gene set (p_corr_ = 0.013), representing 7.9 % of genes included within the GO term (GO:0015833). Functional brain activation in cortical regions yielded no significant overlap with biological processes.

**Fig. 4:**
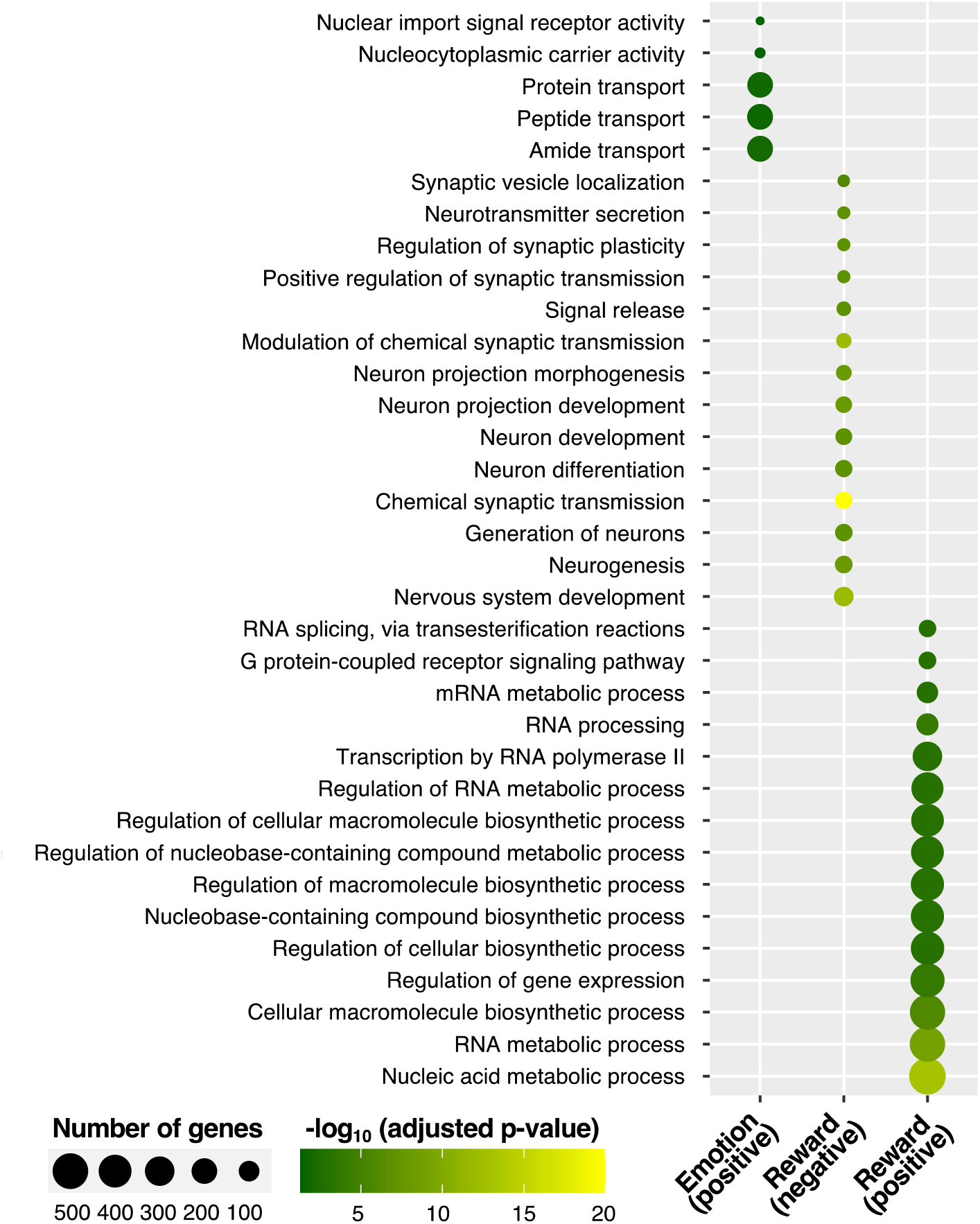
Enriched biological programs for emotion and reward processing based on ontological structure. Specific biological processes (y-axis) are significantly associated with imaging data in the subcortex. The dot size represents the number of genes that are listed within a particular gene ontology (GO) group. Considering the multitude of possible associations of 18,686 genes with all listed GO terms, analyses were restricted to higher ontological hierarchies (GO levels above 4). Only reasonable correlations between imaging and gene expression data were analyzed, with a cut-off value of rho < −0.5 and > 0.5, respectively. Considering genes with an expression pattern positively correlated with emotion processing, 5 biological programs yielded significance, primarily including cellular transport processes. Besides 203 genes associated with the GO term peptide transport, we found an overlap of 199 genes related to protein transport (GO:0015031, p_corr_ = 0.017, 7.89 %), 205 genes involved in amide transport (GO:0042886; p_corr_ = 0.023, 7.82 %), 10 genes associated with nucleocytoplasmic carrier activity (GO:0140142; p_corr_ = 0.041, 28.57 %), as well as 7 genes related to nuclear import signal receptor activity (GO:0061608; p_corr_ < 0.05, 41.18 %).

Analyzing neural responses during acceptance of monetary rewards in subcortical regions, associated GO terms mainly included genes with positive mRNA-fMRI correlations, predominantly comprising transcription processes. Highest significance was present for nucleic acid metabolic process (GO:0090304, p_corr_ < 0.001, 536 genes, 8.95 %), RNA metabolic process (GO:0016070, p_corr_ < 0.001, 486 genes, 8.89 %), as well as cellular macromolecule biosynthetic process (GO:0034645, p_corr_ < 0.001, 485 genes, 8.45 %). GO terms associated with genes negatively correlated with the acceptance of monetary rewards within subcortical regions mainly related to synaptic processes and neuronal development. Thereby, most significant GO terms were chemical synaptic transmission (GO:0007268, p_corr_ < 0.001, 54 genes, representing 6.19 % of genes included within this term) and modulation of chemical synaptic transmission (GO:0050804, p_corr_ < 0.001, 35 genes, 6.4 %). However, in the cortex only two overlapping GO terms were found for genes positively associated with reward processing (autophagy of mitochondrion, GO:0000422, p_corr_ < 0.05, 4 genes, 3.96 % and sensory perception of sound, GO:0007605, p_corr_ = 0.04, 4 genes, 2.34 %).

Focusing on the reward system, altogether 78 and 27 biological processes were associated with genes that correlated negatively and positively with imaging data, respectively (for illustrative purposes, depicted GO terms are limited to the most significant associations).

### 3.4 The role of risk genes implicated in depression

The relationship between neuronal activation patterns and 42 functional genes associated with MDD obtained from a pre-defined gene set was investigated to evaluate the superordinate role of these risk genes on specific BOLD activation. Single-site master regulator analysis of co-regulatory networks based on previously compiled ranked lists including strongest mRNA-fMRI correlations and revealed individual candidate master regulators for each paradigm that potentially regulate genes spatially associated with imaging data. We found 4 regulators from the MDD risk gene set for emotion and 3 for reward processing in subcortical regions, with 79 and 89 possible targets, respectively (p < 0.001) (Table 3).

**Table 3:**
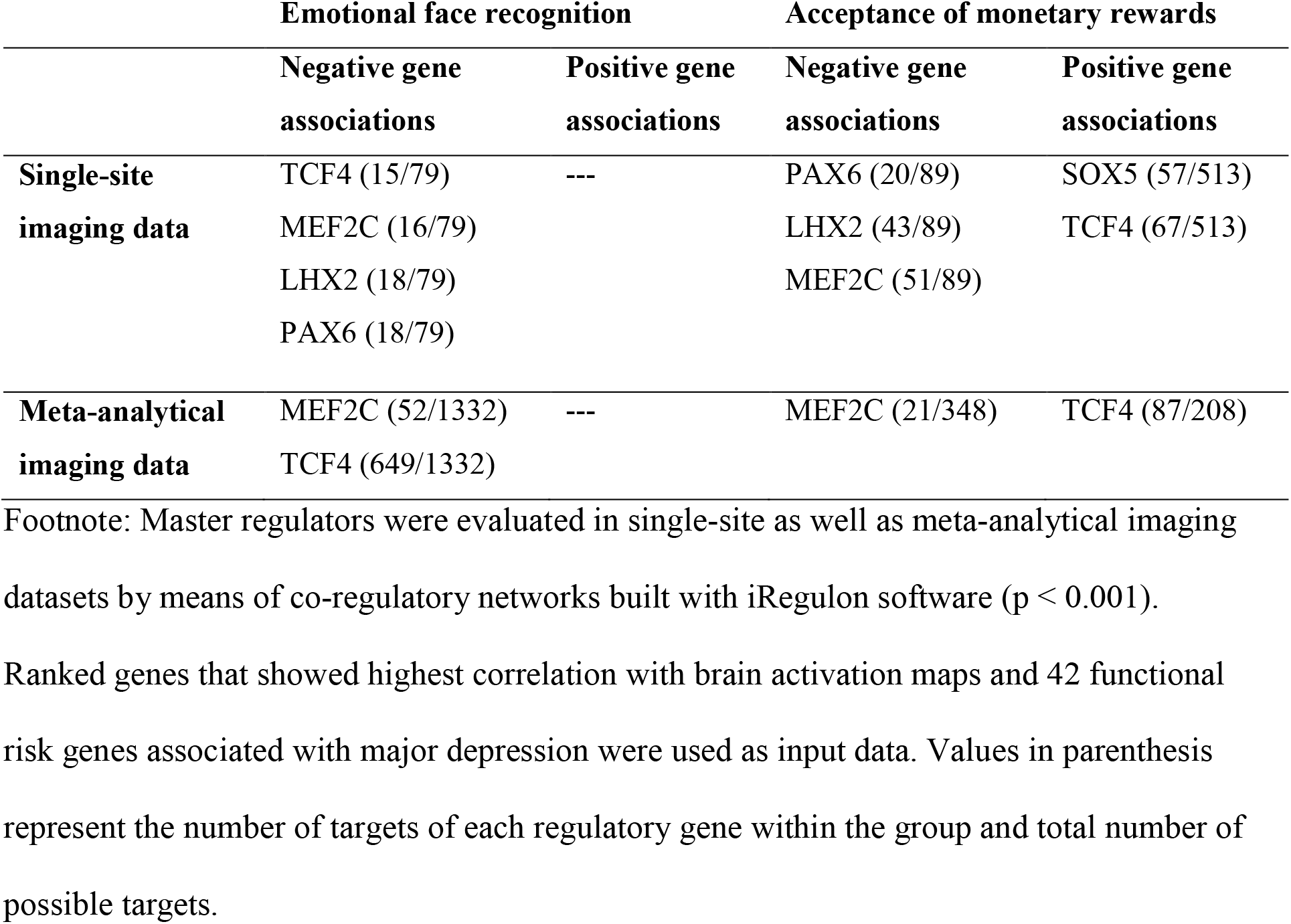
Master regulator analysis in subcortical regions for emotion and reward processing.

Curiously, PAX6, LHX2, as well as MEF2C were associated with negatively correlated genes for both paradigms, indicating a rather superordinate role of these master regulators in MDD, regardless of cognitive system. Alternatively, TCF4 was identified both as a regulator for genes negatively correlated with emotion as well as genes positively correlated with reward processing. Overall, strongest regulation was found for genes negatively correlated with reward processing, whereby LHX2 and MEF2C coordinated more than half of the possible targets. Considering regulation for genes that showed positive spatial correlations with brain activation patterns, two master regulators (SOX5 and TCF4) were identified for reward processing, while no significant regulators were found for genes positively correlated with emotion processing. While corresponding cortical analyses yielded no significant results, subcortical results were validated by means of an independent meta-analytical dataset from the Neurosynth framework. Identically, MEF2C emerged as regulator of genes negatively correlated with emotional face recognition as well as acceptance of monetary rewards (Fig. 5A), while TCF4 showed an inversed regulation of genes correlated with the emotion (negative association) and reward (positive association) paradigm (Fig. 5B).

**Fig. 5:**
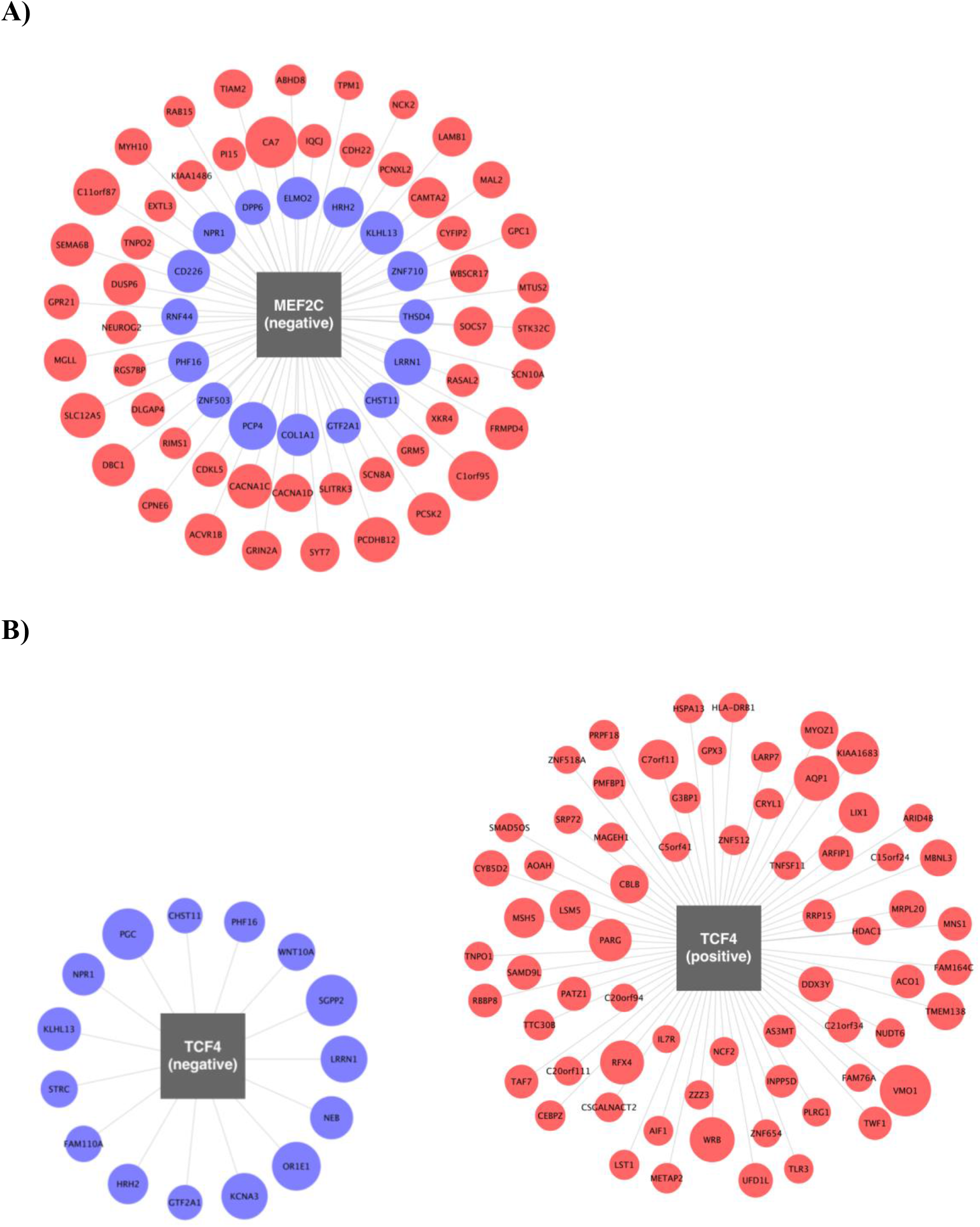
Relationship between risk genes for depressive disorders and genes correlated with functional brain imaging. Two master regulators associated with depressive disorders and corresponding subordinate genes correlated with emotion and reward processing present in single-site and meta-analytical data sets are presented. Graphical visualizations are based on associations between gene expression and single-site imaging data in subcortical regions (p < 0.001). The size of each circle corresponds to the absolute value of Spearman’s correlation coefficient of the respective gene. **A)** MEF2C mainly regulates genes with expression patterns negatively correlated with emotion (16 targets, blue) and reward processing (51 targets, red). **B)** In contrast, TCF4 inversely regulates genes showing negative associations with emotion (15 targets), but positive associations with reward processing (67 targets).

Both, single-site and meta-analytical results from GSEA complemented our findings from GO as well as master regulator analyses, showing an inversed aggregation of MDD risk genes for emotion and reward processing. Rather than focusing solely on ranks of single genes, by means of GSEA we assessed the role of the whole gene set associated with depressive disorders. Regarding emotion processing, risk genes were predominantly enriched within positively correlated genes (maximum ES: 2.881, p = 0.43) in single-site data, albeit not reaching statistical significance. In contrast, an enrichment of risk genes was present within negative correlations for reward processing (maximum ES: −0.2, p = 0.527). Within the Neurosynth dataset, the maximum subcortical ES yielded 0.275 (p = 0.051) for emotion and −0.243 for reward processing (p = 0.65), respectively (Fig. 6). In the cortex, GSEA yielded identical enrichment of MDD risk genes compared to the subcortical analyses (Supplementary Fig. 8).

**Fig. 6:**
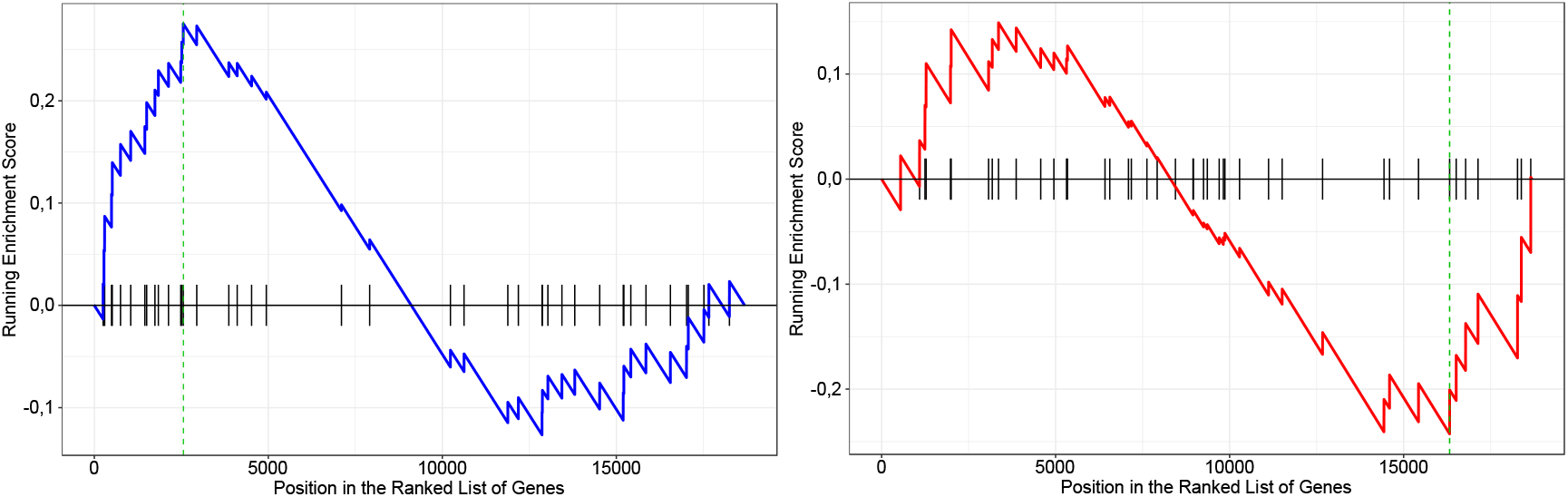
Gene Set Enrichment Analysis (GSEA) for emotion and reward processing, including risk genes implicated in major depression. The analysis revealed an inversed relationship between emotion and reward processing in the subcortex. Vertical lines on the x-axis represent positions of 42 functional risk genes within each ranked list including 18,686 genes, dashed lines mark the locations of the maximum enrichment score (ES) which draws a density line depicting the peak enrichment of risk genes. GSEA with data from the Neurosynth framework yielded a positive value for emotion (ES = 0.275, p = 0.051; blue graph) and a negative value for reward processing (−0.243, p = 0.65; red graph).

## 4 Discussion

### 4.1 Integration of human gene expression and functional imaging data

Here, we applied a comprehensive and integrative methodological approach to investigate the relationship between regional gene expression patterns and macroscopic BOLD responses elicited by emotional face recognition and the acceptance of monetary rewards, under the assumption that strongly correlated genes would coincide with distinct biological programs. Large-scale screening for spatial associations between mRNA expression and functional brain activation resulted in ranked lists of 18,686 genes positively and negatively correlated with BOLD signaling in healthy male and female subjects. Similar distributions of Spearman’s correlation coefficients were present for emotion (ranging from rho = −0.739 to rho = 0.865 in subcortical and from rho = −0.449 to rho = 0.431 in cortical areas) and reward processing (from rho = −0.788 to rho = 0.81 in the subcortex and from rho = −0.639 to rho = 0.698 in the cortex). Exploring the GO knowledgebase, we detected a task-specific overrepresentation of processes related to cellular transport as well as neuronal development, synapse regulation, and transcription for emotion and reward processing, respectively. The enriched ontological terms were interrelated within each fMRI paradigm, thus indicating unique associations of biological programs with different RDoC domains. Focusing on systems for social processes and positive valence systems, a meta-analytic validation sample including over 90 fMRI studies was obtained from the Neurosynth framework. In contrast to a previous study that identified gene-cognition associations based on the Neurosynth framework (Fox et al., 2014), we increased spatial resolution and advanced probe selection of gene expression data by application of interpolated mRNA maps presented by Gryglewski et al. (2018). Given the relevance of imaging paradigms examining subthreshold depressive symptoms within the general population (Lewinsohn et al., 2000), MDD risk genes were eventually analyzed with regard to physiological brain activation. We identified master regulators associated with depressive disorders and task-specific functional brain activation, TCF4 and MEF2C, in two independent datasets. MEF2C primarily regulated genes showing negative correlations with emotion and reward processing. In contrast, TCF4 appeared simultaneously as a regulator for genes negatively correlated with the emotion task, but positively correlated with reward processing. Noteworthy, analyses including meta-analytic brain activation maps eventually confirmed our findings originating from single-site fMRI measurements. Albeit not reaching statistical significance, the supplementary genome-wide association studies (GWAS) enrichment analysis suggested an inversed distribution of genes implicated in major depression. MDD risk genes showed a rather positive association with imaging data for emotional face recognition but a negative association for the acceptance of monetary rewards.

### 4.2 Implications for imaging transcriptomics and depressive disorders

Regarding imaging genetics studies, common candidate genes associated with increased risk for neuropsychiatric disorders may exert distinct effects on brain structure or function (Rose & Donohoe, 2013). Since numerous individual and environmental factors interact with potential genotype effects, sufficiently powered sample sizes are required to detect a significant impact on neuroimaging correlates. However, imaging genetics studies commonly neglect transcriptional and post-transcriptional mechanisms that impact on the actual expression of disorder-related genes (Maričić et al., 2020). Genetic influences on emotional face recognition or adaptive reward-based decision-making have only been evaluated for the mere presence of single gene variants of functional proteins, irrespective of their topological distribution across the brain (Rose & Donohoe, 2013; Wolf et al., 2015). By including whole-brain gene expression patterns, our approach can be discriminated from previous imaging genetics studies investigating effects of individual disease-related single-nucleotide polymorphisms or environmental factors. This study adds up to numerous wide-ranging research findings combining topological mRNA expression with fMRI measures (Richiardi et al., 2015; Shin et al., 2018) or other neuroimaging properties, partially offering toolboxes for an integrative data analysis (Rizzo et al., 2016; Unterholzner et al., 2020). Furthermore, the hierarchically structured GO database was successfully applied in an attempt to find a relationship between well-defined biological processes and imaging properties. The strong interrelation of GO terms within each imaging paradigm certainly illuminate the role of gene expression patterns for human brain functioning (Ashburner et al., 2000).

Most notably, topological expression patterns of designated genes implicated in major depression and their impact on neuroimaging properties have not yet been investigated. Particularly in MDD, additive genetic effects may attribute to individual differences in the phenotype and highlight the importance of large-scale data in systems medicine to resolve unsettled genetic influences on neuroimaging paradigms (Fabbri et al., 2017). While over 322 million people worldwide suffer from depressive disorders, a number that increased by 18.4 % between 2005 and 2015 (James et al., 2018), a significant part of the population is also affected by subthreshold depressive symptoms, potentially originating from different levels of genetic susceptibility in relevant neuronal systems. Our findings endorse the characterization of neuropsychiatric disorders in terms of functions rather than diagnoses, which was recently highlighted within the much-noticed RDoC framework (Sanislow et al., 2019). In line with the debilitating symptoms of depressive disorders, we investigated paradigms reflecting principal functions of both the reward responsiveness construct within the positive valence systems domain as well as the social communication construct, which is part of the systems for social processes domain. Hyper-as well as hypoactivations of brain regions involved in the integration of social information together with a reduced reward sensitivity suggest a polygenic nature of depressive symptoms with distinct imaging features (Ghaemi & Vohringer, 2011; Knutson et al., 2008; Luijten et al., 2017). We endorse the evaluation of physiological brain activation for integrative analyses of post-mortem gene expression and in vivo imaging data. The direct comparison of brain activation maps from depressed individuals or disorder-related maps from the Neurosynth framework with data from the AHBA appears less beneficial, especially since pathological brain activation patterns measured with task-based fMRI affect only few individual brain regions. Given the data disparity between whole-brain gene expression and functional activation maps including only the brain regions that are altered in depressed individuals, the comparison with physiological brain activation during psychological processes appears more reasonable. Nonetheless, identifying a core set of risk genes implicated in subthreshold neuropsychiatric disorders is complicated due to widespread and disease-specific network interactions. A potential solution to this key challenge in systems biology might be the analysis of master regulating genes as well as their corresponding regulons (Janky et al., 2014). Hereof, we present two transcription factors that regulate downstream networks formed by genes strongly correlated with imaging data for two major domains of brain functioning.

### 4.3 Molecular mechanisms associated with neuroimaging properties

Overall, regulating genes associated with MDD included protein coding MEF2C (Myocyte Enhancer Factor 2C), which plays a role in neuronal development, as well as hippocampus-dependent learning and memory (Barbosa et al., 2008; Paciorkowski et al., 2013). Relevance for synapse regulation arises due to alternatively spliced transcript variants involved in neuronal processes, e.g. activated TLR4 signaling or the cAMP response element-binding protein (CREB) pathway. Besides depressive disorders, associations of this transcription activator with other psychiatric disorders like schizophrenia (Mitchell et al., 2017) or attention-deficit/hyperactivity disorder (Shadrin et al., 2018) have been reported. Most notably, significant dependence of striatal neuronal activation was described for an identified risk variant in the TMEM161B-MEF2C gene cluster during a reward task, endorsing well-known deficits of reward processing in MDD observed as anhedonia (Muench et al., 2018). Besides, we affirm previously reported relationships between neuropsychiatric disorders and mutations of TCF4, which has been implicated not only in depression, but also in schizophrenia and autism (Amare et al., 2019; Li et al., 2019). The transcription factor is mainly characterized by its regulatory role for the proliferation and differentiation of neuronal and glial progenitor cells (Ross et al., 2003). Interestingly, the master regulatory role of TCF4 in schizophrenia was recently demonstrated by Torshizi et al. (2019), based on the analysis of transcriptional networks in two independent datasets.

Since the master regulators reported in this study exhibit their effects by numerous molecular mechanisms, a closer investigation of the genes strongly associated with imaging parameters will prospectively allow elaborated statements about regional protein biosynthesis and the allocation of resulting proteins to cellular compartments. For example, DNA-binding transcription factor FOXN4 (Forkhead Box N4), belonging to the Forkhead-box (FOX) superfamily, showed highest correlation with the emotional face recognition paradigm in the cortex. FOX transcription factors are involved in regulatory biological processes and mutations in forkhead genes have been linked to developmental disorders in humans due to substitutions or frameshifts that disable or remove the DNA binding domain (Carlsson & Mahlapuu, 2002). The subtype FOXN4 thereby expresses developmental functions in neural and non-neural tissues, particularly during spinal neurogenesis by modulating a specific expression mosaic of other proneural factors (Misra et al., 2014). Further relevance for neural development was shown by Chen et al. (2014), who demonstrated the location of FOXN4 on neurons and astrocytes as well as an increased expression after spinal cord injury lesions. Although associations with depressive or other neuropsychiatric disorders have not been published, the role of FOXN4 as a key transcriptional regulator during developmental processes demands further research, especially since the full set of its targets in the CNS are not known yet. Accordingly, the C10orf125 gene (Fucose Mutarotase, FUOM), expressed in the brain and other tissues, showed highest correlation with emotional face recognition in the subcortex. The corresponding gene transcript, fucose mutarotase, is an enzyme of the fucose-utilization pathway performing the interconversion between α-L-fucose and β-L-fucose on human cell surfaces. Hereof, besides one animal study demonstrating male-like sexual behavior in FUOM knock-out mice, presumably resulting from reduced fucosylation during neurodevelopment (Park et al., 2010), further associations with pathological states in mammals have not been published for this gene. Regarding reward processing, the protein coding DUSP3 (Dual-specificity phosphatase 3) gene, member of the dual-specificity protein phosphatase subfamily, showed strongest correlation with measured fMRI data in the cortex. Members of these protein tyrosine phosphatases (PTPs) regulate the phosphorylation of the mitogen-activated protein (MAP) kinase signaling pathway and control cell signaling, especially in regard to cytoskeleton reorganization, apoptosis and RNA metabolism (Tonks, 2013). DUSP3 shows a wide expression in different tissues as an opposing factor of protein tyrosine kinases (PTKs) and acts as a central mediator of cellular proliferation and differentiation. Whereas a role in neoplastic disorders, pathologies related to immunology, angiogenesis as well as Parkinson’s disease (PD) have been related to anomalous tyrosine phosphorylation (Cohen & Alessi, 2013; Russo et al., 2018), associations with psychiatric disorders have not been described. In subcortical regions, MDK (midkine) was among the highest correlating genes for the reward paradigm. MDK transcribes one of two growth factors from the heparin-binding cytokine family and plays a role during differentiation of neurons, especially within dopaminergic pathways (Alguacil & Herradón, 2015). Notably, it facilitates neuroprotective effects in neurodegenerative disorders, drug-induced neurotoxicity in the striatum, or after neural injury (Yoshida et al., 2014). Disease-related publications suggest accumulation of midkine in senile plaques and increased serum levels in patients with Alzheimer’s disease (Salama et al., 2005), genetic variations associated with PD (Prediger et al., 2011), and an influence on addictive behaviors (Gramage et al., 2013). While a possible therapeutic target of MDK has already been proposed for autoimmune diseases including multiple sclerosis (Takeuchi, 2014), more recently, elevated serum levels of this neurotrophic factor were also associated with autism spectrum disorder (Esnafoglu & Cirrik, 2018).

### 4.4 Limitations of the study

Results of this study are limited due to the structure of applied data sets and reflect rather subtle genetic influences on psychological processes, disregarding the dynamic nature of short-term regulatory mechanisms, environmental factors, or individual variations due to genetic ancestry. Since a whole-brain proteome atlas has not been published yet, spatial integration of functional imaging data relies on publicly available mRNA maps. While gene expression levels don’t necessarily reflect actual in vivo protein densities (Komorowski et al., 2020), signal alterations in fMRI studies are also impaired by confounding variables, such as network-architectures, structural and functional connectivity measures, or non-specific brain activation. To minimize BOLD signaling elicited by superimposed executive functions, control conditions need to be implemented for each paradigm, which ensures specificity of the performed tasks. Although the inclusion of meta-analytical maps minimizes a potential bias originating from the smaller population size of the single-site data, an exclusive utilization of the Neurosynth framework would reduce the functional specificity due to rather broadly defined terms of interest that lack control conditions. Overall, the congruence of the results appears indeed valid, given fact that both datasets don’t correlate completely. Lower mRNA-fMRI correlations in the cortex compared to the subcortex may be partially ascribed to data analysis in volume space, which is a less precise representation of the cortical sheet than the surface space (Coalson et al., 2018). However, considering strongest brain activation elicited by emotion processing in ventral striatum, amygdala, ventral tegmental area, fusiform gyri, insula, and medial prefrontal cortex, in that order, it seems plausible that higher correlation levels were observed in subcortical regions. Regarding the reward paradigm, functional activation was likewise more prominent in subcortical brain regions compared to the cortex (Goya-Maldonado et al., 2015). The insignificant results of the performed GSEA were partly hampered by the small number of the risk genes implicated in depression. The enrichment of the risk gene set was further affected by the occurrence of master regulators modulating up- and downregulation of subordinate risk and non-risk genes. Still, the GWAS performed by Wray et al. (2018) is among the largest ever conducted in psychiatric genetics and provides a solid basis for further research about the genetic architecture of MDD.

Studies investigating post mortem gene expression are principally limited due to locally and functionally regulated epigenetic and epitranscriptomic modifications that affect actual protein distribution (Maier et al., 2009). Inter-individual differences regarding effects of age or genotype cannot be considered, when performing integrative analyses on the basis of the AHBA that originally derived expression values from only 6 post-mortem brains (Hawrylycz et al., 2012; Shen et al., 2012). Since female gene expression was acquired from solely one donor, sex-specific differences also have to be neglected. Besides outdated annotation information and inaccurate sample assignment, ubiquitous noise due to expression of genes with a low spatial dependence hampers data analysis. Although interpolated whole-brain gene expression data suffer from spatial autocorrelation that may inflate correlations analyses with any relatively smooth fMRI data, voxel-wise results were corroborated in the region-wise analyses in this study. In general, these issues have been thoroughly addressed by Gryglewski et al. (2018) during the creation of the transcriptome maps, which allow for seamless integration with neuroimaging modalities.

Despite advances and decreased costs of high-throughput gene expression profiling, the necessity for large cohorts in genetic studies calls for collaborations and integrative approaches. Within the framework of future studies, the vast potential of the AHBA might even be increased by re-assigning available mRNA probes to corresponding genes on the basis of the latest sequencing information to increase the number of specifically annotated genes. Harmonized data processing pipelines or methodological guidelines instead of rather unique approaches to data integration and corresponding statistical measures could further increase comparability between studies (Arnatkevičiūtė et al., 2019; Keil et al., 2018; Kim et al., 2014).

### 4.5 Conclusion

This investigation highlights the advantages of a comprehensive approach to reveal genetic influences on functional brain imaging by integrating imaging and large-scale transcriptome data with sufficient power. While traditional imaging genomics strategies have solely investigated the effects of individual genotype variations, the analysis of whole-brain gene expression patterns provide superior information for the understanding of neuropsychiatric disorders. In line with recent insights from the RDoC framework, this multimodal study highlights the relevance of functional brain activation related to social interaction and the experience of reward for systems medicine. Both, the emotion and the reward system are associated with specific biological programs like cellular transport, neuronal development, synapse regulation, and transcription processes. The identification of regulatory genes TCF4 and MEF2C thereby implies a potential role of master regulating genes for functional brain imaging. This work exemplifies an integrative approach including complementary information from multiscale data, which seems to be increasingly relevant in the big data era.

## Acknowledgements

We thank the students of the Laboratory of Systems Neuroscience and Imaging in Psychiatry (SNIP-Lab) and the Neuroimaging Labs (NIL) for support. Parts of this study have been presented by AK at the 32nd ECNP Congress, 07-10 September 2019, Copenhagen, Denmark. With relevance to this work there is no conflict of interest to declare. AS and RGM are funded by the German Federal Ministry of Education and Research (Bundesministerium fuer Bildung und Forschung, BMBF: 01 ZX 1507, ‘‘PreNeSt - e:Med’’). MM is funded by the Austrian Science Fund FWF DOC 33-B27. GG was recipient of a DOC-fellowship of the Austrian Academy of Sciences at the Department of Psychiatry and Psychotherapy, MUV. JW is supported by an Ilídio Pinho Professorship, iBiMED (UID/BIM/04501/2013) and FCT project PTDC/DTP-PIC/5587/2014 at the University of Aveiro, Portugal.

## Author contributions

RGM and RL conceived the study; AK and RGM designed the study, primarily interpreted results, and drafted the article; RGM, RV, AS, MM, TPC, and GG made significant contributions to data collection, including quality control, data processing and statistical analysis; SK and JW revised the study and contributed to the intellectual content. All authors have critically revised the article and approved it for publication.

## Data availability

The functional imaging data generated during the current study are available from the corresponding author upon reasonable request. The correlation lists generated in this study, the associated biological processes, as well as the analyzed risk genes are included in the supplementary information files. All meta-analytical imaging datasets analyzed are available in the Neurosynth repository, https://neurosynth.org/, whole-brain transcriptome maps are available from the Neuroimaging Labs (NIL), www.meduniwien.ac.at/neuroimaging/mRNA.html, and gene ontology data are available from the Gene Ontology Consortium, http://geneontology.org/.

## Code availability

No custom algorithm or software that are central to the conclusion have been used in this study. The codes that were included in this study are available from the corresponding authors upon reasonable request. To process data, licensed software as well as freely accessible webbased tools were used, whereby the authors presume further availability without restrictions on existing code or algorithm availability in the foreseeable future.

