## Supplementary material for "Regional gene expression patterns are associated with task-specific brain activation during reward and emotion processing measured with functional MRI": Suppl Material

### Supplementary figures

**A)**

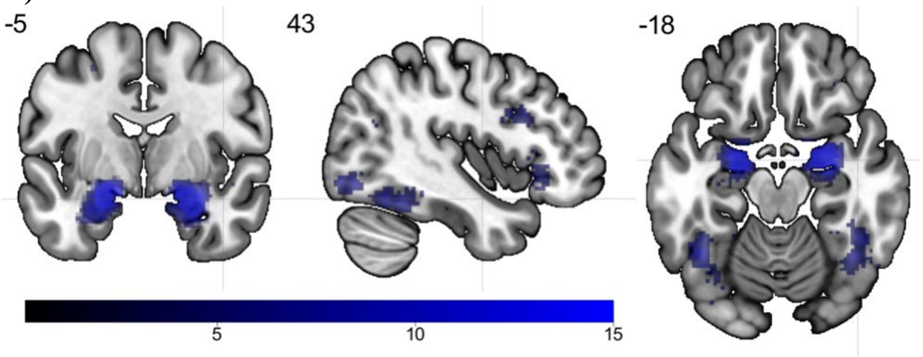


**B)**

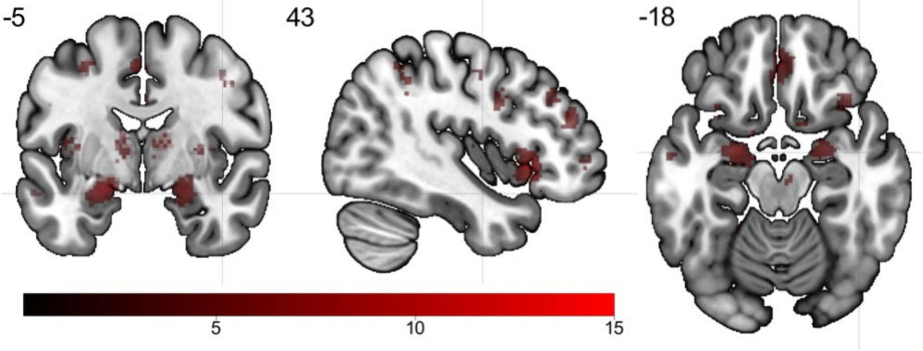


Supplementary Fig. 1: Meta-analytical functional magnetic resonance imaging data (z-score) obtained from the Neurosynth database is visualized in MNI space (activation maps are thresholded above 0 for visualization purposes only). **A)** The uniformity test map related to the term "fearful faces" matched with single-site fMRI data measured during recognition of negative faces. **B)** The uniformity test map related to the term "rewards" matched with single-site fMRI data measured during the acceptance of monetary rewards.

**A)**

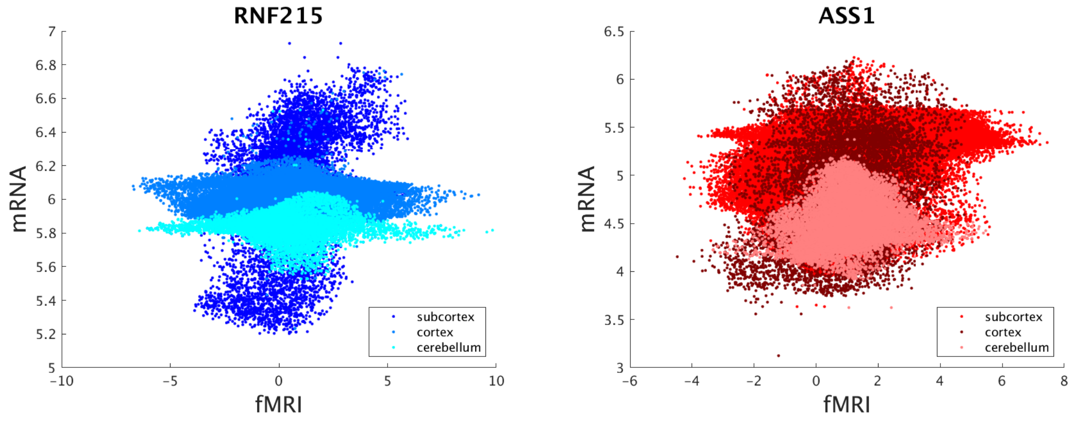


**B)**

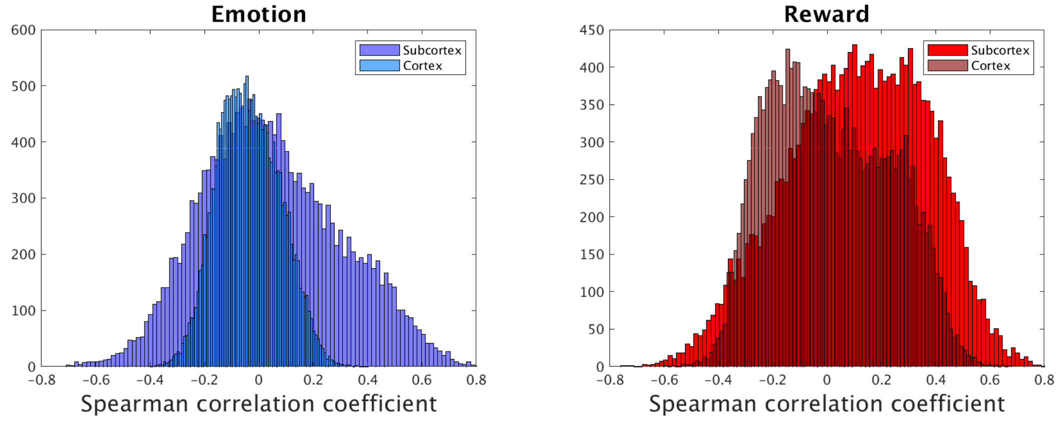


Supplementary Fig. 2: Gene expression differences in cortical, subcortical, and cerebellar structures for emotion and reward processing. **A)** The scatter plots depict voxel-wise correlations (subcortex: 10,863 voxels, cortex: 129,817 voxels, cerebellum: 24,415 voxels) between whole-brain transcriptome maps and single-site imaging data for emotional face recognition (RNF215) and reward processing (ASS1). **B)** Histograms show distributions of correlation coefficients of 18,686 genes for region-wise analyses using the Brainnetome atlas for emotional face recognition and reward processing. Markedly differing expression levels justified separate analyses for each brain structure.

**A)**

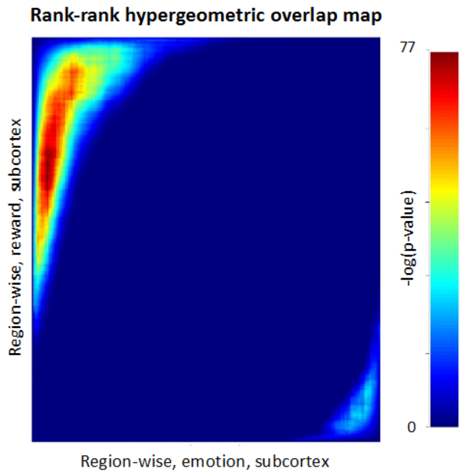


**B)**

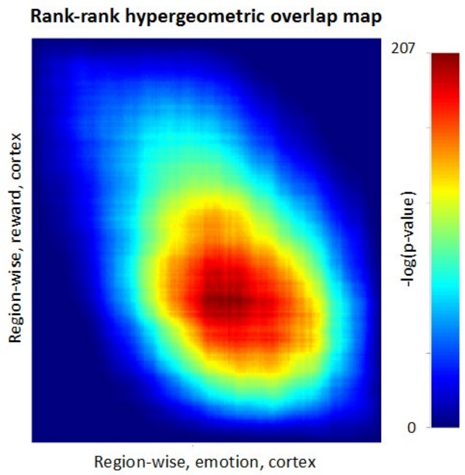


Supplementary Fig. 3: Rank–rank hypergeometric overlap (RRHO) visual representation of single-site imaging data for reward vs. emotion processing. Genes with low agreement of correlation coefficients between both lists (either positive or negative) show lower statistical significance in the bottom left and top right corner. Region-wise RRHO comparing ranked lists including 18,686 genes indicated low congruence between both paradigms in **A)** subcortical ($rho_{RRHO}$ = -0.282) and **B)** cortical structures ($rho_{RRHO}$ = 0.205).

**A)**

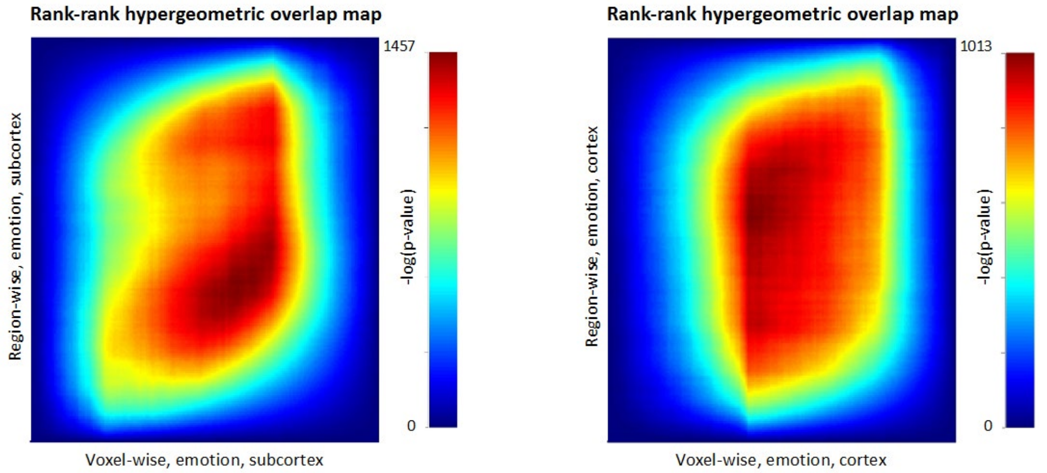


**B)**

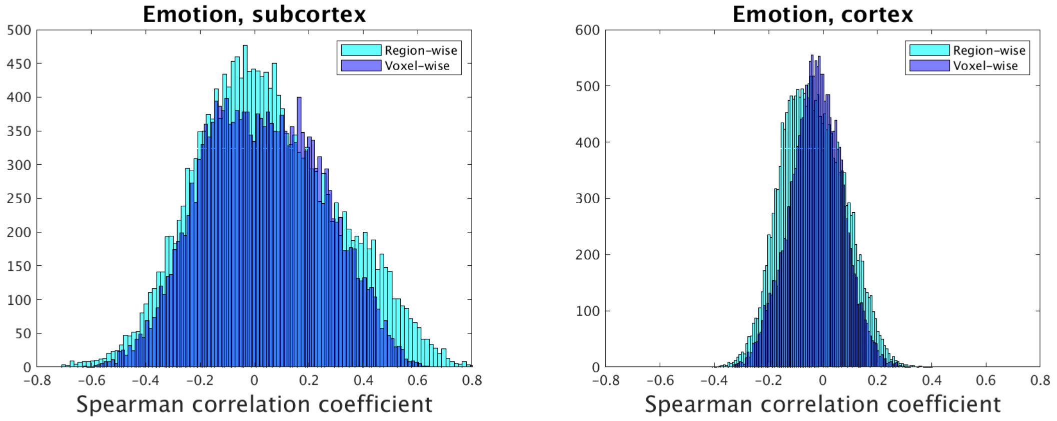


Supplementary Fig. 4: Voxel-wise vs. region-wise correlation analyses of single-site imaging data during emotion processing. **A)** Agreement between compiled lists including 18,686 genes was compared by means of rank–rank hypergeometric overlap (RRHO), which indicated a fairly high congruence for the emotional face recognition paradigm. Visual representations of RRHO depict significance of overlap between ranked lists (warmer colors correspond to lower p-values), comparing the voxel-wise vs. region-wise approach for the subcortex ($\text{rho}_{\text{RRHO}}$ = 0.722) and the cortex ($\text{rho}_{\text{RRHO}}$ = 0.612). **B)** Histograms of Spearman's correlation coefficients applying a voxel-wise as well as a region-wise approach are provided for subcortical and cortical regions.

**A)**

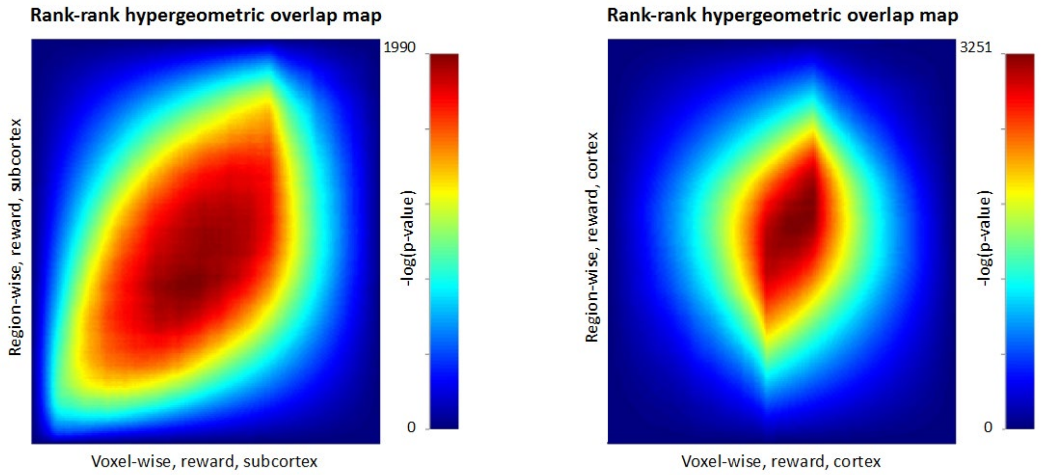


**B)**

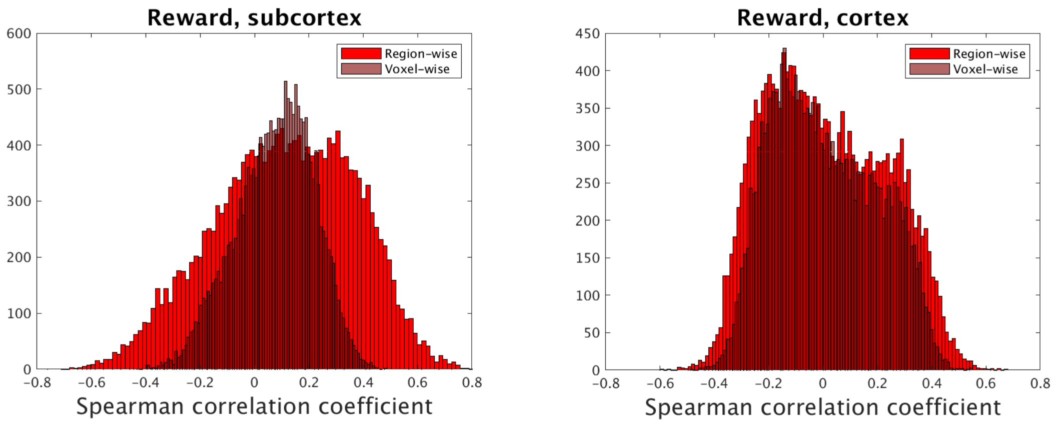


Supplementary Fig. 5: Voxel-wise vs. region-wise correlation analyses for single-site imaging data during reward processing. **A)** Agreement between compiled lists including 18,686 genes was compared by means of rank–rank hypergeometric overlap (RRHO), which indicated a fairly high congruence for the reward paradigm. Visual representations of RRHO depict significance of overlap between ranked lists (warmer colors correspond to lower p-values), comparing the voxel-wise vs. region-wise approach for subcortex ($rho_{RRHO}$ = 0.829) and cortex ($rho_{RRHO}$ = 0.793). **B)** Histograms of Spearman's correlation coefficients applying a voxel-wise as well as a region-wise approach are provided for subcortical and cortical regions.

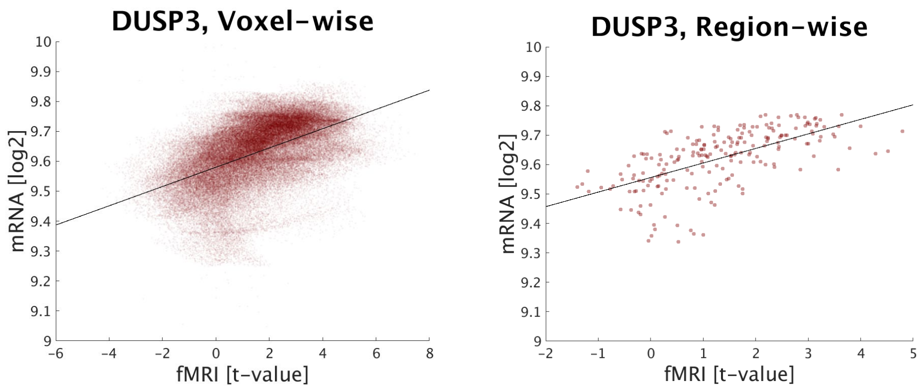


Supplementary Fig. 6: Comparison of functional brain activation during reward processing and mRNA expression of DUSP3 in cortical regions. The scatter plots depict correlations between whole-brain transcriptome maps and single-site imaging data (acceptance of monetary rewards) for voxel-wise (rho = 0.548; 129,817 voxels) and region-wise (rho = 0.698; 210 regions, $p < 0.0001$) analyses. Each dot represents expression values and corresponding imaging parameters at target coordinates or within anatomical regions, respectively.

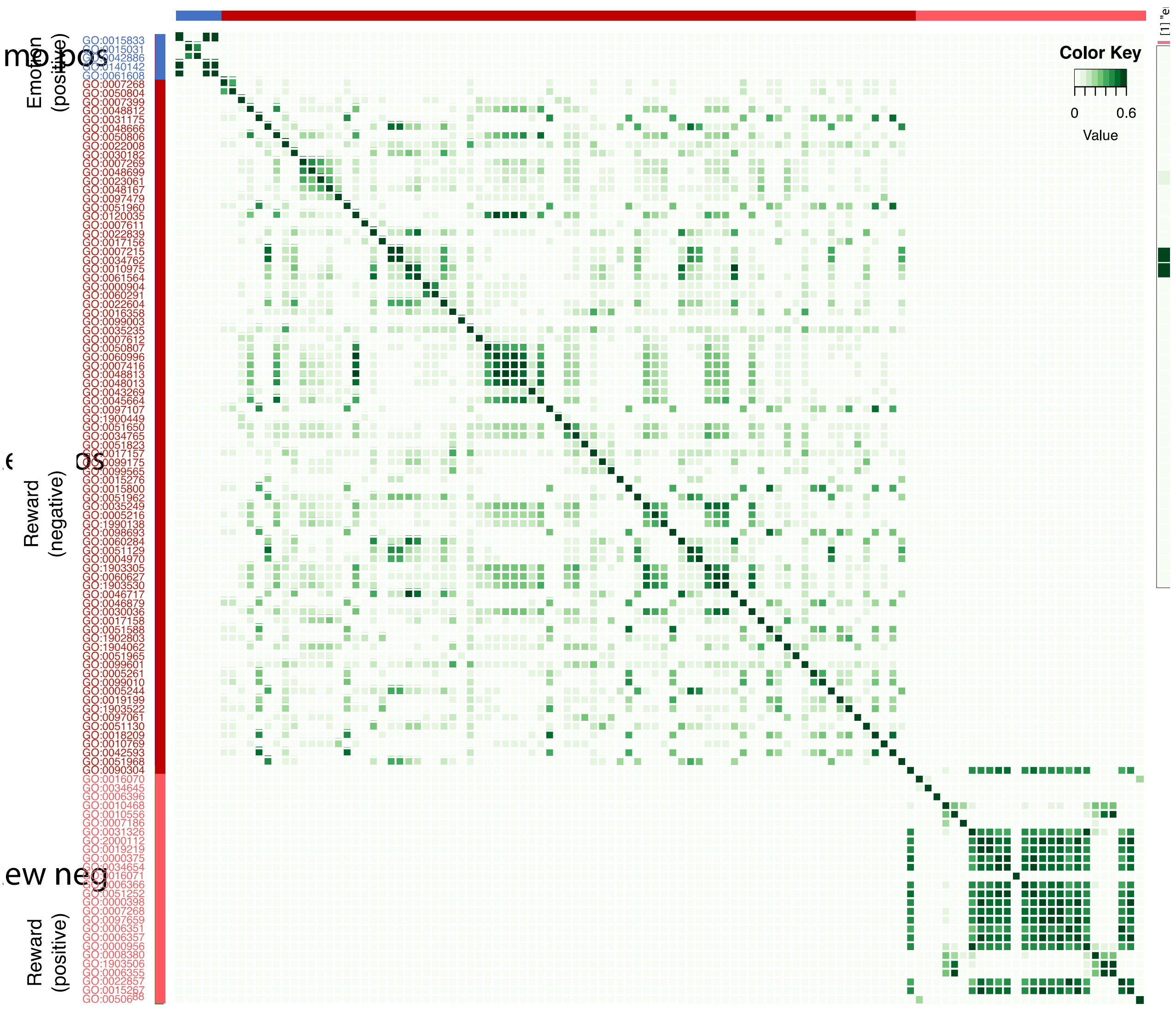


Supplementary Fig. 7: Enriched biological programs for emotion and reward processing based on ontological structure. Potential gene overlaps between imaging conditions are negligible, indicated by low Jaccard indices referring to the significantly enriched biological processes (emotional face recognition: blue; acceptance of monetary rewards: red)

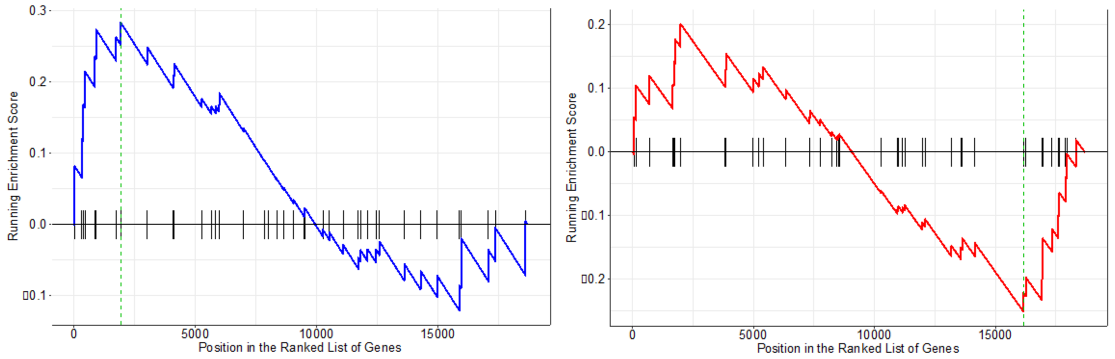


Supplementary Fig. 8: Gene Set Enrichment Analysis (GSEA) for emotion and reward processing, including risk genes implicated in major depression. Vertical lines on the x-axis represent positions of 42 functional risk genes within each ranked list including 18,686 genes; dashed lines mark the locations of the maximum enrichment score (ES) which draws a density line depicting the peak enrichment of risk genes. Analyzing single-site data, GSEA showed an inversed relationship within cortical structures, yielding maximum ES for emotion processing of 0.284 ($p = 0.2$, blue graph) and -0.25 for reward processing ($p = 0.276$, red graph), respectively.

**Supplementary tables**

Supplementary Table 1: Previously published risk genes associated with major depression. The gene set composed of 69 published functional and non-functional risk genes; bold names correspond to 42 functional genes that were included for gene set enrichment and master regulator analyses.

| | | | | |
|---|---|---|---|---|
| **RERE** | **MLF1** | **ASTN2** | **LRFN5** | CRYBA1 |
| **SLC45A1** | **SLC30A9** | **DENND1A** | **SYNE2** | **MYO18A** |
| **NEGR1** | **LINC00682** | **LHX2** | MIR548H1 | **NUFIP2** |
| LINC01360 | DCAF4L1 | **SORCS3** | ESR2 | MIR924HG |
| **DENND1B** | LINC00461 | DKFZp686K1684 | DLST | **DCC** |
| **VRK2** | **MEF2C** | PAUPAR | PROX2 | MIR4528 |
| LINC01876 | LOC101927421 | **ELP4** | **RPS6KL1** | **RAB27B** |
| **NR4A2** | TENM2 | **PAX6** | **BAG5** | **CCDC68** |
| **GPD2** | **FBXL4** | **SOX5** | **APOPT1** | **TCF4** |
| TOPAZ1 | C6orf168 | **ENOX1** | RBFOX1 | MIR4529 |
| TCAIM | **TMEM106B** | LACC1 | SHISA9 | **L3MBTL2** |
| **ZNF445** | **VWDE** | **CCDC122** | **CPPED1** | EP300-AS1 |
| **RSRC1** | PUM3 | **OLFM4** | **PMFBP1** | **CHADL** |
| LOC100996447 | LINC01231 | LINC01065 | **DHX38** | |

Supplementary Table 2: Regions showing functional brain activation during emotional face recognition and acceptance of monetary rewards. Single-site emotion (sad > object) and reward (reward > attention) contrasts are reported at a collection threshold p < 0.001 with k > 10.

| | **BA** | **Region** | **Cluster size (k)** | **Peak voxel (T-value)** | **MNI coordinates** | | |
|---|---|---|---|---|---|---|---|
| | | | | | **x** | **y** | **z** |
| **Emotion contrast** | A37lv_R | Fusiform gyrus right | 1759 | 9.83 | 38 | -38 | -26 |
| | A39c_R | Inferior parietal lobule right | | 7.63 | 42 | -60 | 6 |
| | iOccG_R | Lateral occipital cortex right | | 7.25 | 40 | -72 | -8 |
| | A37lv_L | Fusiform gyrus left | 782 | 7.65 | -40 | -42 | -24 |
| | V5/MT+_L | Lateral occipital cortex left | | 7.20 | -44 | -74 | -2 |
| | A37dl_L | Middle temporal gyrus left | | 5.92 | -50 | -66 | 10 |
| | mAmyg_R | Amygdala right | 708 | 6.47 | 18 | -2 | -22 |
| | A38m_R | Superior temporal gyrus right | | 6.12 | 28 | 10 | -28 |

| | | | | | | | |
|---|---|---|---|---|---|---|---|
| | mAmyg_R | Amygdala right | | 5.92 | 20 | -8 | -14 |
| | rCunG_R | Medioventral occipital cortex right | 1444 | 5.90 | 2 | -66 | 4 |
| | A31_R | Precuneus right | | 5.70 | 4 | -58 | 18 |
| | rLinG_L | Medioventral occipital cortex left | | 4.97 | -18 | -50 | -4 |
| | Outside atlas | Uncus left | 315 | 5.79 | -26 | 6 | -26 |
| | rHipp_L | Hippocampus left | | 5.06 | -18 | -6 | -20 |
| | rHipp_L | Hippocampus left | | 4.54 | -24 | -20 | -20 |
| | A22r_R | Superior temporal gyrus right | 169 | 5.76 | 46 | -8 | -14 |
| | A21r_R | Middle temporal gyrus right | | 5.64 | 52 | 6 | -22 |
| | A9l_R | Superior frontal gyrus right | 152 | 5.05 | 12 | 54 | 28 |
| | A9l_R | Superior frontal gyrus right | | 4.28 | 12 | 56 | 38 |
| | rpSTS_L | Posterior superior temporal left | 46 | 4.83 | -52 | -42 | 8 |
| | A44d_R | Inferior frontal gyrus right | 388 | 4.78 | 46 | 22 | 22 |
| | IFS_R | Inferior frontal gyrus right | | 4.65 | 48 | 32 | 12 |
| | A6cvl_R | Precentral gyrus right | | 4.51 | 42 | 8 | 24 |
| | Outside atlas | Precuneus left | 43 | 4.63 | -10 | -46 | 44 |
| | cpSTS_L | Posterior superior temporal left | 18 | 4.02 | -58 | -50 | 12 |
| | Outside atlas | Parahippocampal gyrus right | 24 | 3.96 | 16 | -42 | 2 |
| | A13_L | Orbital gyrus left | 11 | 3.76 | -2 | 28 | -18 |
| | A38m_R | Superior temporal gyrus right | 21 | 3.74 | 36 | 24 | -36 |
| | A9l_L | Superior frontal gyrus left | 14 | 3.65 | -8 | 54 | 36 |
| | rLinG_R | Medioventral occipital cortex right | 11 | 3.64 | 12 | -50 | -6 |
| **Reward contrast** | Outside atlas | --- | 34203 | 10.85 | -18 | -6 | 50 |
| | A8m_L | Superior frontal gyrus left | | 10.14 | -6 | 6 | 50 |
| | A6cdl_L | Precentral gyrus left | | 10.05 | -42 | -8 | 58 |
| | Outside atlas | --- | 9940 | 10.80 | 30 | -46 | -24 |
| | Outside atlas | Cerebelum left | | 9.34 | -34 | -56 | -24 |
| | Outside atlas | Cerebelum right | | 8.78 | 28 | -60 | -22 |
| | Outside atlas | Frontal middle orbital left | 132 | 5.08 | -24 | 52 | -18 |
| | Outside atlas | --- | 60 | 4.50 | 0 | -30 | 10 |
| | Outside atlas | Middle occipital gyrus right | 10 | 4.29 | 26 | -84 | 14 |
| | A11l_R | Orbital gyrus right | 96 | 4.15 | 22 | 48 | -16 |
| | A10l_R | Middle frontal gyrus right | | 3.94 | 22 | 60 | -14 |
| | A10l_R | Middle frontal gyrus right | | 3.69 | 36 | 56 | -16 |
| | Outside atlas | --- | 21 | 3.88 | 42 | -24 | -8 |
| | Outside atlas | --- | 15 | 3.84 | 64 | 14 | -16 |

Footnote: Anatomical brain regions are labeled according to the Brainnetome atlas (BA).

Supplementary Table 3: Associations between functional imaging and transcriptome data. Correlation analyses yielded comparable results between single-site measurements and the Neurosynth uniformity maps “fearful faces” as well as “rewards”. For both data sets, ranked genes with corresponding Spearman’s correlation coefficients are reported separately in cortical as well as subcortical brain regions; p-values derived from permutation tests are provided for region-wise correlations.

(excel-file)

Supplementary Table 4: Enriched biological programs for emotion and reward processing based on ontological structure. Specific gene sets listed within the gene ontology (GO) knowledgebase significantly overlapped with genes showing expression patterns strongly correlated with single-site imaging data in the subcortex. Cortical associations yielded no meaningful findings. Resulting GO IDs, levels, and corresponding p-values are provided for both paradigms.

(excel-file)
